# Systematic cross-study assessment of RNA-Seq experimental workflows for plasma cell-free transcriptome profiling

**DOI:** 10.1101/2025.07.10.664092

**Authors:** Cristina Tuñí-Domínguez, Giovanni Asole, Pablo Monteagudo-Mesas, Elena Cristina Rusu, Lluc Cabús, Lucila González, Laura Sánchez, Beatriz Neto, Phil Sanders, Marc Weber, Julien Lagarde

## Abstract

Plasma cell-free RNA (cfRNA) is a promising source of non-invasive biomarkers, but its clinical translation is hindered by technical challenges and a lack of protocol standardization, which compromises reproducibility and comparability across studies. There is a need for a systematic evaluation of existing cfRNA-Seq workflows to understand the drivers of technical variability. Here, we address this gap by performing a comprehensive cross-study analysis of 2,166 cfRNA-Seq samples from 15 published studies and an in-house generated dataset, applying a uniform bioinformatics pipeline to enable a controlled comparison of experimental workflows. Our analysis reveals that the donor phenotype typically explains a negligible fraction of the transcriptomic variation, whose main determinants are technical – principally protocol choice, genomic DNA contamination levels and library diversity. Remarkably, this technical noise is so profound that variation within plasma cfRNA samples exceeds that found across a wide range of human tissues. Finally, we demonstrate that critical pre-analytical factors are often confounded with patient phenotypes, jeopardizing the validity of biomarker discovery efforts. Our work serves as a comprehensive benchmark of current cfRNA-Seq methodologies and provides evidence-based guidelines to improve experimental design. By highlighting the dominance of controllable technical factors, we offer a path towards more robust and reproducible cfRNA research.

## Introduction

Plasma cell-free RNA (cfRNA) has recently emerged as a non-invasive analyte of considerable promise, offering a dynamic and systemic view into human physiological and pathological states^1–6^. The circulating cell-free transcriptome, comprising a diverse range of RNA species including messenger RNAs (mRNAs), ribosomal RNAs (rRNAs), long noncoding RNAs (lncRNAs), microRNAs (miRNAs) and transfer RNAs (tRNAs), is released into the bloodstream from various tissues and cell types. Far from being mere cellular debris, this collection of transcription products reflects active biological processes, providing real-time information about cellular function and dysfunction throughout the body. The rich mosaic of cfRNA biotypes holds significant potential for diverse clinical applications, including the early detection of diseases, diagnostic assessment, prognostic stratification, and the monitoring of therapeutic responses across a spectrum of conditions. For example, studies have demonstrated significant disruptions in cfRNA profiles in patients with various cancers^2–4,7,8^, and cfRNAs detected in maternal plasma can predict gestational age and risks of preterm birth or preeclampsia^9,10^.

The diagnostic potential of the plasma cell-free transcriptome extends far beyond the identification and profiling of individual biomarkers. Instead, the whole cfRNA landscape can be conceptualized as a systemic indicator, providing insights into complex biological processes and the overall health status of an individual. Different RNA species perform distinct biological functions, and their respective abundance in circulation can paint a detailed picture of the body’s response to disease. The capacity to comprehensively and accurately profile this diverse array of cfRNA molecules is therefore critical. Transcriptome-wide sequencing (RNA-Seq) is the method of choice for this purpose, as it enables the discovery of novel biomarkers and offers a holistic view of gene expression changes.

However, despite its immense potential, technical barriers have so far prevented the translation of cfRNA-Seq into the clinic. A primary hurdle is the biochemical instability of RNA in solution, which leads to exceedingly low concentrations of cfRNA in plasma, combined with high levels of molecular fragmentation^11^. cfRNA is susceptible to degradation by ubiquitous ribonucleases (RNases) present in blood and surfaces, necessitating meticulous sample handling and processing. Each step of the cfRNA-Seq workflow (from blood extraction to bioinformatics analysis) represents a critical control point where variability, bias and error can be introduced, potentially compromising the integrity and interpretation of the final results^11–13^.

The expansion of the cfRNA research field has occurred in the relative absence of comprehensive, systematic evaluations that compare the performance of different cfRNA-Seq workflow methodologies and their effects on data quality and biological interpretation. Most studies examining the influence of pre-analytical variables have either been limited to miRNAs, or confined to targeted mRNA analysis using PCR-based methods on a small number of genes^14–17^. This has contributed to a lack of standardization, hindering the reproducibility and comparability of findings across different laboratories. Adding to this challenge, studies often employ varied and non-standard bioinformatics analysis pipelines, which further complicates direct cross-study comparison. Ultimately, this slows the validation of promising cfRNA biomarkers for clinical application, underscoring the urgent need to assess and standardize these workflows.

Here, we present a comprehensive cross-study analysis of 2,166 cfRNA-Seq samples, comprising data from 15 published studies and an in-house generated dataset (Flomics). To provide a comparative baseline, we also include reference datasets from cell-free DNA-Seq (N=90 samples) and bulk tissue RNA-Seq (N=100 samples). Using this large collection, we systematically assess the impact of experimental workflows on data quality and biological interpretation. By applying a uniform bioinformatics pipeline specifically tailored for long RNA analysis (such as mRNA and lncRNA), we perform a controlled comparison of protocol performance and quantify the primary sources of variation across datasets. Our analysis reveals a profound lack of methodological standardization and demonstrates that technical variables – primarily genomic DNA contamination and library diversity – together with inter-laboratory and inter-protocol batch effects, are the dominant drivers of variation, often masking biological signals. Furthermore, we evaluate the impact of non-human sequence contamination on transcriptomic profiles. Using computational deconvolution, we show that the cellular origin of cfRNA is also shaped by pre-analytical factors. Ultimately, this work provides a detailed benchmark of existing protocols and offers a set of evidence-based recommendations to guide the design of more robust and reproducible cfRNA-Seq studies for biomarker discovery.

## Results

### Lack of standardization in pre-analytical and library preparation protocols

We curated a diverse collection of cfRNA-Seq data from 15 published studies^2–4,7–9,18–26^, supplemented by an in-house dataset (Flomics, unpublished) and, for reference, samples from bulk tissue RNA-Seq (ENCODE/EN-TEx^27^) and cell-free DNA-Seq (cfDNA, Wei^28^) (Fig. 1A, see Methods). In total, 2,356 Illumina-sequenced samples were selected for analysis (Supplementary table 1). For clarity, datasets are named after the first author of the corresponding publication. In some cases, datasets were subdivided to reflect distinct sample batches within a study, based on our manual curation of available metadata (see Methods).

**Figure 1.**
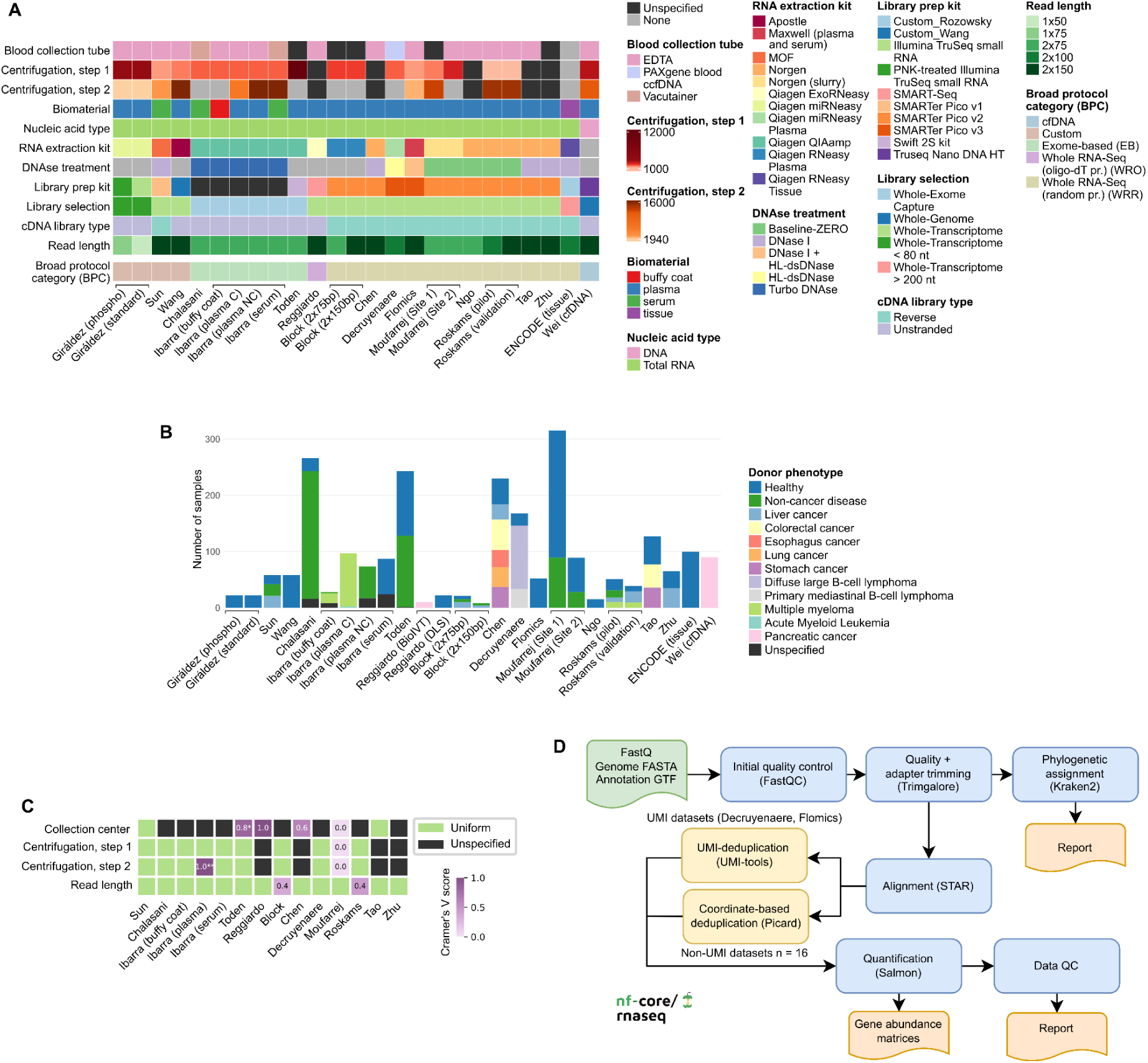
Overview of datasets and processing pipeline. Sample batches identified within some studies are depicted separately but grouped together using horizontal brackets below the X axis. “Unspecified”: not reported in the publication. **(A)** Summary of pre-analytical variables across datasets. Datasets were split into distinct batches when those differed by at least one pre-analytical variable. Note that the Toden dataset is annotated per their published Methods, which do not mention DNase treatment. As discussed later in the main text, we speculate that this likely reflects an unintentional omission from the authors, rather than a genuine absence of treatment. **(B)** Overview of datasets under consideration, broken down by donor phenotype. **(C)** Association between various pre-analytical variables and donor phenotype across datasets. Cramér’s V scores are reported for each variable. Variables strongly confounding phenotype approach a score of 1. “Uniform” means that the variable of interest is constant within the dataset. Only datasets studying more than one phenotypic condition were included. For clarity, we also excluded pre-analytical variables that were fully uniform or unspecified across all datasets. Those were: “Blood collection tube”, “RNA extraction kit”, “DNase treatment”, “Library prep kit”, “Library selection” and “cDNA library type”. *One sample with missing information from the Toden dataset was removed to allow the calculation. **17 samples with missing phenotype information from the Ibarra (plasma) dataset were removed. (**D)** Illustration of the nf-core/rnaseq Nextflow pipeline used for the uniform processing of RNA-seq data.

Our cross-study analysis revealed a profound lack of standardization in experimental protocols, with significant differences observed both across and within individual studies. Pre-analytical steps such as blood centrifugation showed major disparities. Some studies employed a single centrifugation step while others used two, with wide variability in both speed and duration (Fig. 1A). Furthermore, these critical parameters were often insufficiently reported, as described previously^29^, hindering reproducibility.

Similar variability was also observed in the library preparation protocols. While most studies utilized established commercial kits, others employed custom methods. The choice of reverse transcription priming was also inconsistent. In contrast to the random priming used by most protocols, the Reggiardo dataset employed an oligo-dT-based strategy^30^. This latter approach is generally considered unsuitable for cfRNA, as the high degree of RNA degradation means most fragments lack the polyA tail required by this technique. The majority of datasets consisted of whole-transcriptome libraries, with three studies using exome capture instead. Moreover, a number of studies did not perform DNase treatment to remove contaminating genomic DNA (gDNA). To systematically analyze this heterogeneity, we grouped the studies into the following broad protocol categories (BPC) based on their library selection and construction methods: “cfDNA”, “Custom”, “Exome-based” (EB), “Whole RNA-Seq (oligo-dT-primed)” (WRO), “Whole RNA-Seq (random-primed)” (WRR) (Fig. 1A).

### Confounding pre-analytical factors in cfRNA-Seq studies

The datasets included in this analysis exhibited considerable variability in scope (Fig. 1B). While most studies focused on comparisons between healthy and disease states, four included only healthy individuals, and four compared healthy controls to multiple cancer types. The cohorts represented a wide range of conditions, including healthy controls (N=979), various cancers (N=739), and non-cancer diseases (N=572).

Within datasets that included samples from healthy and disease subjects, we observed that the phenotype was sometimes confounded with pre-analytical variables (Fig. 1C). In particular, the collection center was partially confounded with the phenotype in the Chen (Crámer’s V score = 0.6) and Toden (score = 0.8) studies, and was completely confounded in the Reggiardo study (score = 1.0), where pancreatic cancer and healthy samples originated from two different providers. This raises the concern that the observed differences in cfRNA-Seq profile could stem from variations in sample processing between providers rather than the disease state itself. Other confounding factors included read length (Block and Roskams-Hieter, score = 0.4) and centrifugation protocol (Ibarra (plasma), score = 1.0), which were partially and fully confounded with the phenotype, respectively. In contrast, the Moufarrej dataset demonstrated appropriate experimental design by evenly distributing control and disease samples across its two collection sites and centrifugation protocols (score=0.0), thereby minimizing the risk of confounding the results. Importantly, assessing the full impact of these variables was hindered by the lack of metadata reporting. For example, information about the collection center was not specified in 8 of the 14 datasets, making it impossible to assess its influence in those cases.

### A best-practice, uniform data processing pipeline

To ensure a standardized analysis of all datasets, we processed all samples with a single, uniform bioinformatics pipeline, nf-core/rnaseq^31^ (Fig. 1D). Implemented in the Nextflow^32^ language, the pipeline encapsulates best-practice methodologies for NGS analysis of long RNAs, including extensive data quality assessment, mapping of reads with STAR^33^, and subsequent gene quantification using Salmon^34^. Pipeline run settings were adapted to individual datasets via configuration files according to relevant pre-analytical variables (see Methods and Supplementary table 2).

### Cell-free RNA libraries are enriched in microbial sequences

An initial quality control analysis revealed substantial disparities across all datasets in both sequencing depth (Supplementary fig. 1A) and the proportion of reads mapping to the human genome (Fig. 2A). This variability in mapping efficiency was strongly associated with the BPC employed: cell-free WRR libraries exhibited significantly lower mapping rates compared to those from bulk tissue (ENCODE). In contrast, EB protocols (Chalasani, Ibarra, Toden) yielded high mapping statistics comparable to those of tissue-derived libraries. Notably, a stark divergence was observed even within the same protocol, with the Roskams-Hieter “pilot” and “validation” cohorts showing markedly different median mapping rates of 31% and 87%, respectively, despite an identical processing workflow^3^.

**Figure 2.**
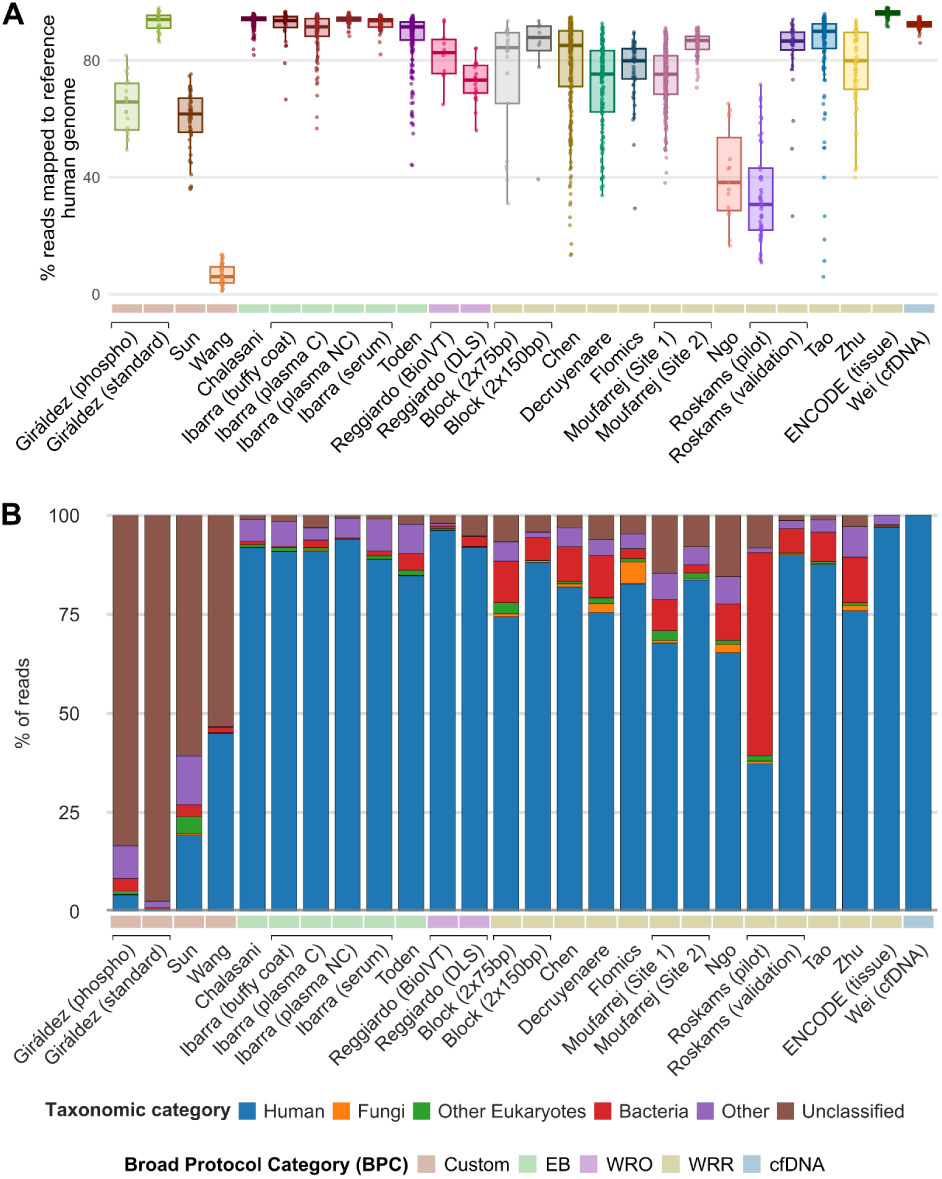
Microbial reads in cfRNA-Seq libraries. **(A)** Fraction of sequencing reads mapping to the human genome across datasets. Each library is represented by a dot. The distribution of the metric within a given dataset is represented by a boxplot. **(B)** Taxonomic profile of dataset batches. Relative abundance values were averaged across samples, grouped into the simplified categories: human (Homo sapiens), fungi (Fungi), other_eukaryotes (Eukaryota except human and Fungi), bacteria (Bacteria), other (Archaea, Viruses, and other unspecific taxa), and unclassified.

We hypothesized that the low mapping efficiency in cfRNA-Seq samples was due to the presence of non-human sequences. To test this, we performed taxonomic classification of raw sequencing data for all samples using the Kraken2 software^35^. This analysis revealed that bulk tissue RNA libraries consisted predominantly of human sequences (97 +/- 4.6% mean relative abundance +/- SD), while cell-free WRR-based samples showed a markedly reduced fraction of human reads (77.4 +/- 20.3%). In these libraries, the non-human fraction was primarily composed of microbial (*i.e.*, fungal or bacterial) sequences (50.2 +/- 26.6% mean +/- SD, excluding Kraken2-unclassified reads, Fig. 2B, Supplementary fig. 2). Hence, we posit that the poor mapping rates in affected samples are largely explained by a high proportion of microbial-derived sequences (Supplementary fig. 3), which fail to align to the human genome due to sequence divergence. EB libraries were largely devoid of microbial reads (2.1 +/- 4.6%), consistent with their use of human-specific capture probesets. Similarly, WRO-based libraries were mainly devoid of microbial reads (2.1 +/- 2.3%, mean microbial fraction +/- SD), consistent with the general lack of compatible polyA tails in bacterial sequences. Custom protocols were instead characterized by an enrichment in sequences unclassified by Kraken2 (median fraction: 83.5%, 60.7% and 53.3%, respectively). This is possibly attributable to the shorter effective fragment length (EFL, which depends on both cDNA fragment size and sequencing fragment length) of these libraries (Supplementary fig. 1B), a factor known to impair the software’s classification accuracy^35^.

The observed differences in microbial sequence enrichment could stem from several sources: the experimental protocol, laboratory contaminations, or genuine biological differences in the cell-free sequence composition of the donors, as reported elsewhere^2,24,36^. In the Roskams-Hieter study, the “pilot” cohort exhibited a significantly higher microbial read fraction (52.0%) compared to its “validation” counterpart (6.4%) (Mann-Whitney U test p=2.8e-15). This disparity strongly suggests varying contamination levels during experimental procedures, as both cohorts included non-cancer and cancer phenotypes, making true biological differences in circulating microbial RNA abundance unlikely. Similar observations apply to the Moufarrej study, which shows differences in microbial fractions between the Site 1 (8.6%) and Site 2 (2.4%) batches (p = 2.5e-26), and different collection centers in the Chen study (Supplementary fig. 2G, PKUFH vs. Southwest Hospital p = 3.1e-10, PKUFH vs. SMMU (EHBH) p = 2.4e-02).

### Genomic DNA contamination: a major pitfall in cell-free RNA profiling

Prior studies have highlighted genomic DNA (gDNA) contamination as a significant challenge in cfRNA analysis^12^. As gDNA fragments lack distinctive features to differentiate them from most RNA-derived reads, they can skew transcriptome profiles and gene quantification^37,38^. We assessed the gDNA contamination level across all datasets using the fraction of spliced reads (FSR) as a primary proxy (Fig. 3A). Since only RNA-derived fragments are expected to span splice junctions, FSR is inversely correlated with the level of gDNA contamination. This analysis revealed substantial variation in gDNA contamination across studies. As expected, cfDNA samples from the Wei study consistently exhibited near-zero FSR (0.08 +/- 0.01%, median +/- SD), confirming the metric’s validity. High levels of gDNA contamination were strongly associated with the absence of DNase treatment during library preparation. For instance, several cfRNA-Seq datasets lacking DNase treatment (Block, Giráldez, Sun and Wang, Fig. 1A) displayed extremely low FSR values (0.80 +/- 2.50%), suggesting these samples consisted almost entirely of gDNA.

**Figure 3.**
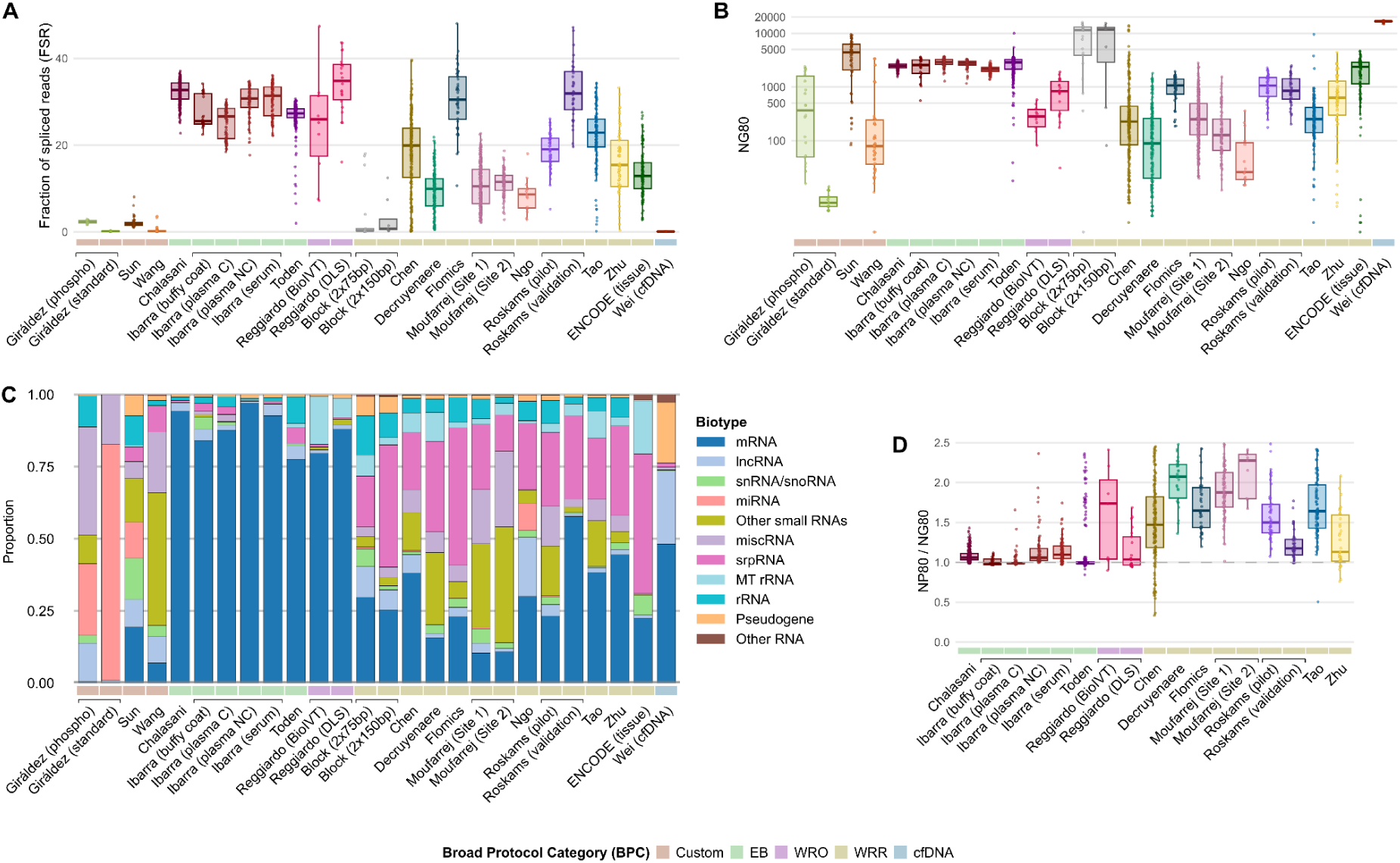
gDNA contamination and library diversity in cfRNA-Seq libraries. **(A)** Fraction of spliced reads. **(B)** Library diversity (NG80). **(C)** Mean RNA biotype representation. **(D)** NP80/NG80 ratio in gDNA-free datasets.

Conversely, EB and WRO datasets showed consistently low levels of gDNA, though the underlying mechanism differs between the two categories. For WRO, oligo-dT priming is genuinely RNA-selective: only polyadenylated molecules are reverse-transcribed, excluding gDNA at the priming step. For EB, this is not due to the capture selection of transcribed sequences, since it selects fragments by complementarity to exonic bait sequences, a criterion met equally by cDNA and gDNA fragments that happen to overlap exonic regions. The low gDNA content of the three EB datasets instead reflects DNase digestion applied upstream of capture: Chalasani *et al.* and Ibarra *et al.* both report DNase treatment prior to reverse transcription (Fig. 1A). Toden *et al.* do not explicitly mention this step, but given the shared lab origin (namely, Molecular Stethoscope Inc.) and near-identical extraction protocol across all three studies, we consider it likely that this study used the same DNAse-based experimental workflow. The low gDNA content of EB samples is therefore attributable to the DNAse treatment step rather than an intrinsic property of the exome capture chemistry.

In addition to protocol-driven differences, we observed significant batch effects within some studies. The two Reggiardo cohorts displayed markedly different median FSR values (24.7% for BioIVT vs 34.1% for DLS, MWW test p=0.011), possibly due to differences in the quality of RNA libraries that were generated. The Roskams-Hieter “pilot” cohort had a much higher gDNA contamination level than its “validation” counterpart (18.8% vs 32.3%, respectively, MWW test p=5.5e-14). This disparity correlated with our previous observation of a higher microbial read fraction in the same “pilot” batch. We hypothesize that the “pilot” cohort consists mainly of significantly degraded RNA. The loss of endogenous RNA fragments would introduce a compositional bias, artificially inflating the relative abundance of more stable contaminants like human gDNA and microbial sequences.

Finally, alternative proxies for gDNA contamination, such as the fraction of exonic reads (FER, Supplementary fig. 1C) and data strandedness (Supplementary fig. 1D), yielded results largely consistent with the FSR analysis. A notable exception was the Decruyenaere dataset, which showed high FER but low FSR. This specific discrepancy may be attributable to an enrichment of genuine, small unspliced transcripts in these libraries, as reported in the following section. Based on these metrics, we classified dataset batches with a median FSR > 10% or FER > 75% as “gDNA-free”. The studies meeting these criteria (at both the dataset and batch level) were Chalasani, Chen, Decruyenaere, Flomics, Ibarra, Moufarrej, Reggiardo, Roskams-Hieter, Tao, Toden, and Zhu.

### Impact of protocol on cfRNA library diversity

Effective biomarker discovery from cfRNA-Seq requires a comprehensive and informative representation of the circulating transcriptome’s landscape. In this context, library diversity (LD) serves as a critical quality metric, providing a faithful proxy to a library’s information content. To quantify and compare LD across studies, we employed the NG80 metric – the number of genes accounting for 80% of a sequencing library’s reads. A higher NG80 value indicates greater LD, reflecting a more even capture of the cfRNA profile without domination by a few highly abundant transcripts.

Our analysis revealed marked heterogeneity in LD across datasets (Fig. 3B). All EB enrichment protocols produced consistently diverse libraries (2,571 +/- 656, median NG80 +/- SD), comparable to those from bulk tissue RNA (ENCODE, 2,375 +/- 1,122). In contrast, gDNA-contaminated datasets (Block, Sun) displayed high NG80 values (11,545 +/- 5,497 and 4,429 +/- 2,766, respectively). This is likely an artifact of uniform genomic sampling from contaminating gDNA, which mimics the high LD observed in pure cfDNA libraries (Wei, 16,680 +/- 423). gDNA-free whole-cfRNA-Seq studies (including both WRO and WRR datasets) consistently displayed lower library diversities (261 +/- 1,023). Across the analyzed datasets, NG80 exhibited a strong relationship with established metrics of diversity and inequality such as Shannon entropy (*R*=0.73, Supplementary fig. 4A) and Gini index (*R*=-1, Supplementary fig. 4B). NG80 offers important advantages over these measures: it is mathematically bounded and provides a more conceptually intuitive representation of transcriptomic complexity.

We hypothesized that the variability in LD across BPCs may be linked to different protocols enriching for or selecting against specific gene biotypes. To investigate this, we analyzed the composition of each library by aggregating gene-level relative abundances (transcripts per million, TPM, see Methods) into predefined gene categories (Fig. 3C). This analysis revealed that the library’s biotype profile was fundamentally dictated by the BPC methodology, particularly the underlying capture and reverse transcriptase (RT) priming strategy. As expected, EB datasets yielded libraries composed predominantly of protein-coding genes (mRNA, 91.8 +/- 14.7% median fraction +/- SD), a direct consequence of their targeted probe design. In stark contrast, WRR datasets had a lower fraction of protein-coding fragments (21.0 +/- 17.9%), and were instead comparatively enriched in various noncoding RNAs. Signal recognition particle RNAs (srpRNAs) were particularly prominent in this category, whereas they were detected at negligible levels in capture-based libraries, suggesting their exclusion from the capture probe sets. Across these diverse strategies, ribosomal RNA (rRNA) depletion was consistently effective, with rRNA constituting only a minor fraction of most libraries (2.42 +/- 6.51%).

The importance of the library construction method was further underscored in the WRO-based samples where libraries rich in protein-coding genes were produced. This likely occurred due to the use of oligo-dT RT priming, which selectively enriches poly-adenylated transcripts. This selectivity explains the depletion of non-polyadenylated genes such as srpRNAs in these libraries. The high srpRNA fraction observed in the ENCODE bulk tissue data (mean: 48.32 +/- 22.99%) is explained by its reliance on random priming, the same method used by the WRR cfRNA protocols.

To confirm that the lower LD in WRR-based datasets is driven by abundant non-coding RNAs, we calculated the LD of the coding transcriptome, the NP80 metric, defined as the number of genes accounting for 80% of a sequencing library’s *protein-coding* reads. We then computed the ratio of NP80 to the overall LD (NG80) for each sample in gDNA-free datasets (Fig. 3D). A NP80/NG80 ratio approaching one indicates that LD is primarily driven by protein-coding genes, whereas a higher ratio suggests that abundant non-coding transcripts are reducing the overall NG80 value. As predicted, capture-based datasets exhibited a ratio close to one (1.03 +/- 3.17, median NP80/NG80 +/- SD). WRR datasets, in contrast, displayed a substantially higher ratio (2.68 +/- 40.90), with the most pronounced increases in datasets dominated by non-coding RNAs (Decruyenaere, Flomics, Moufarrej). These results indicate that the reduced LD in whole-transcriptome datasets is a direct consequence of the overwhelming abundance of specific non-coding RNA subclasses, which are otherwise depleted by EB or WRO-based methods.

### The cellular origin of cfRNA is mainly hematopoietic and shaped by pre-analytical variables

While plasma cfRNA reflects contributions from all blood-irrigated tissues and thus holds great diagnostic potential, its composition can be biased by experimental factors leading to cellular RNA contamination, as reported previously^1,11^. To investigate how these experimental variables shape the cellular contributions to the cell-free transcriptome, we performed computational deconvolution on all gDNA-free cfRNA-Seq datasets to predict the relative abundance of contributing cell types. Across all studies, the cfRNA landscape was dominated by only six major cell types (MCTs, see Methods) (platelets, monocytes, erythrocytes, B cells, macrophages and neutrophils), all of them hematopoietic, consistent with previous findings^5,39^. These six MCTs contributed, on average, 69.28% of the cell-free transcriptome (SD: 12.74%, Fig. 4A and Supplementary fig. 5), with erythrocytes and platelets being the two most abundant cell types across all samples (56.62 +/- 16.89%, average fraction +/- SD).

**Figure 4.**
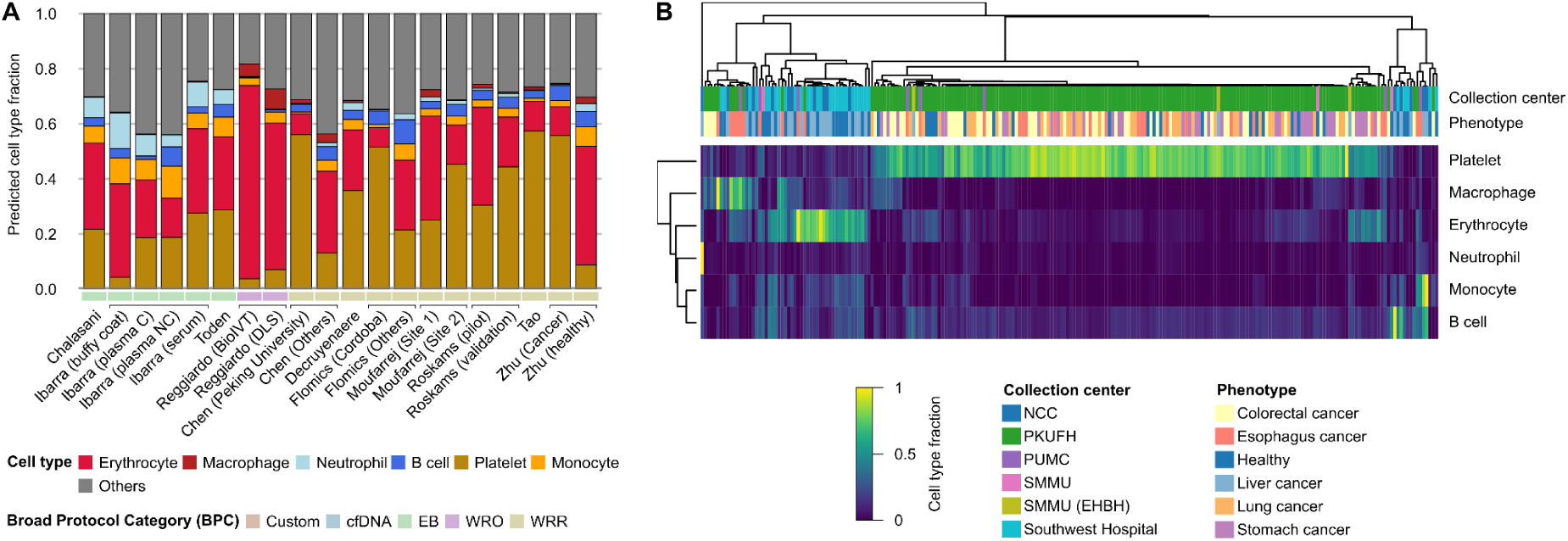
Cellular origin of the cell-free transcriptome. Only gDNA-free datasets are represented **(A)** Average predicted cell type contribution to cfRNA. **(B)** Clustered heatmap showing a cell type fraction batch effect in the Chen dataset. Only major cell types (MCTs) are represented on the Y axis. Heatmap cells are colored according to the corresponding predicted cell type contribution to the cfRNA library (X axis), in relative abundance. Each library is annotated with its corresponding relevant metadata (“phenotype” and “collection_center”, top rows)

Clustering samples based on relative MCT contribution revealed that inter-sample heterogeneity was primarily driven by pre-analytical variables rather than donor phenotype (Supplementary fig. 6). For instance, distinct sample clusters in the Chen (Fig. 4B) and Flomics (Supplementary fig. 6A) datasets were strongly associated with specific collection centers. Similarly, differences in centrifugation protocols (double vs single centrifugation) between the two Moufarrej collection sites led to a significant disparity in the contribution of platelet-derived RNA (Supplementary Fig. 6D). This finding is consistent with recent studies linking platelet RNA contamination with centrifugation settings in cfRNA-Seq data^1^. Furthermore, the two Reggiardo batches exhibited high-erythrocyte signatures consistent with high levels of hemolysis during plasma extraction (Fig. 4A).

Notably, the Zhu dataset was the only case where samples clustered clearly by phenotype (healthy vs liver cancer), with the clustering being driven by a large difference in platelet fraction between the two groups (8.9% in healthy vs 55.8% in liver cancer, MWU test p=7.4e-12). While the magnitude of this difference in platelet RNA representation raises the strong possibility of a technical batch effect, we cannot conclusively rule out a genuine biological signal without orthogonal validation.

### Determinants of cfRNA profile variation are predominantly technical

A primary goal of cfRNA-Seq is the discovery of disease-related gene signatures^1–6^. Therefore, we investigated and quantified the influence of unwanted technical variables on gene expression variation across all 2,356 samples in our study using dimensionality reduction techniques – principal component analysis (PCA) and t-distributed stochastic neighbor embedding (t-SNE). PCA revealed that the first two components (PC) captured 68.35% of the total variation (42.89% for PC1 and 22.46% for PC2, Fig. 5A). When visualized, samples separated into distinct groups that corresponded to their original studies (Fig. 5A) and their associated BPC (Figure 5B), an effect even more evident in t-SNE maps (Supplementary fig. 7A). Of note, datasets from the same laboratory (Chen, Tao, Zhu) clustered together. A targeted sub-analysis restricted to healthy controls (N=979) demonstrated that samples clustered consistently by laboratory and protocol, confirming that technical batch effects remain the dominant source of variation even within a biologically more homogeneous cohort (Supplementary fig 7B). Collectively, these results highlight the major influence of experimental protocols on the cfRNA profiles observed.

**Figure 5.**
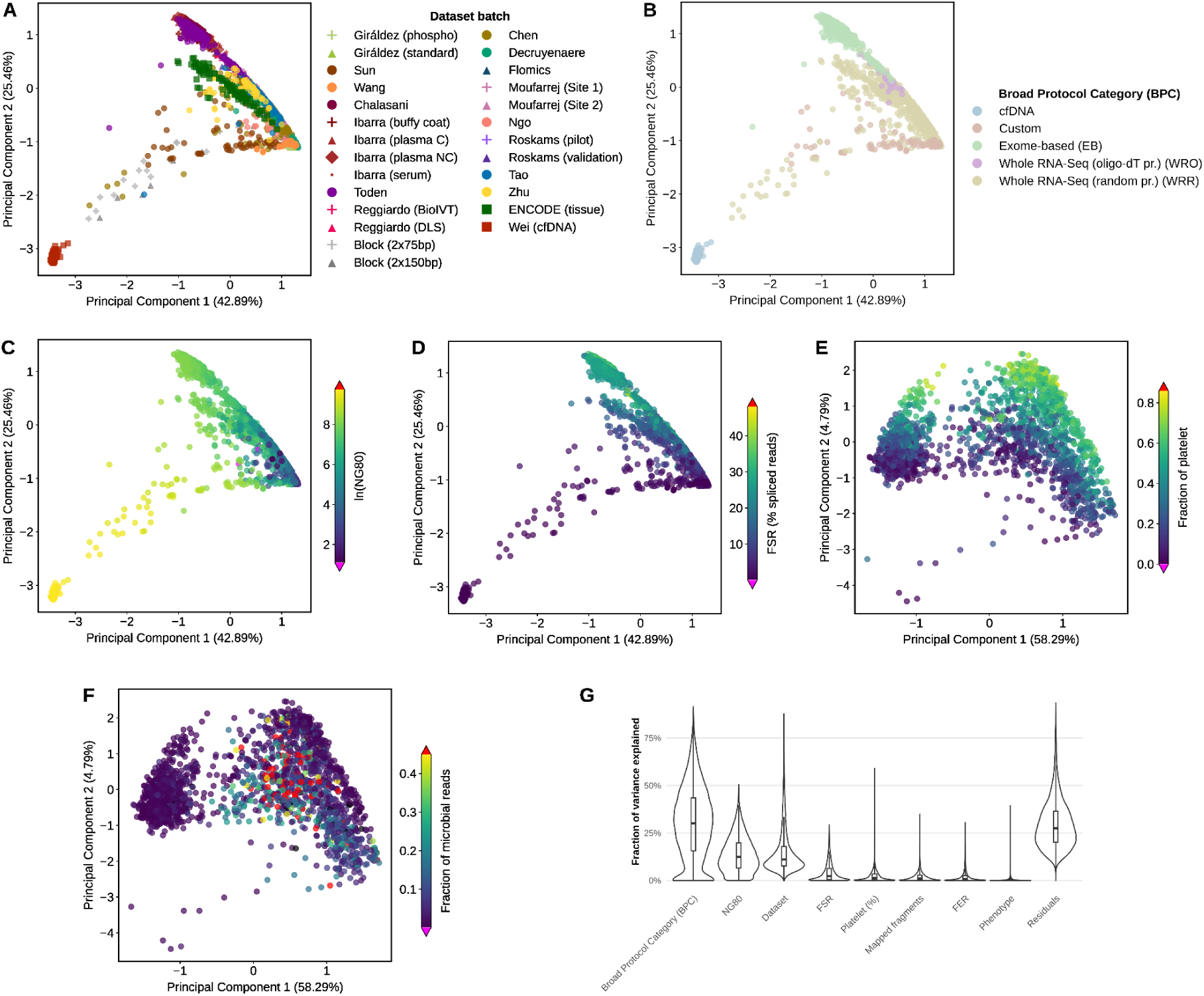
Determinants of cell-free RNA gene abundance profiles. **(A-F)** Scatterplot visualizations of gene abundance-based principal components 1 and 2. Each dot corresponds to a sequencing library. **(A-C)**: All datasets. **(E-F)** gDNA-free datasets only. In this latter representation, note that the left and right clusters roughly correspond to EB and WRR/WRO datasets, respectively (Supplementary fig. 7I). **(A)** Colored by dataset/batch label. **(B)** Colored by BPC (Broad Protocol Category). **(C)** Colored by log(NG80). **(D)** Colored by fraction of spliced reads (FSR). **(E)** Colored by fraction of platelet RNA. **(F)** Colored by fraction of microbial reads. **(G)** Variance Partition Analysis (VPA). For each variable of interest (X axis), a genome-wide violin plot of the distribution of variance explained by this variable across all genes is shown. Variables are sorted in descending order by median variance explained across genes, except the residuals, which are represented on the right by convention.

Consistent with this protocol-driven separation, the primary driver of cell-free transcriptome variation was library diversity (measured by NG80, Fig. 5C), which was strongly correlated with PC1. The second major driver was the gDNA contamination level (measured by FSR, Fig. 5D), which was strongly correlated with PC2 and thus has a strong influence on the cfRNA profile. Given that library diversity is itself mainly driven by technical variables such as gDNA contamination levels and the library preparation strategy, these findings confirm the dominance of technical factors. As expected, cfDNA samples were strong outliers in the PCA. Attempts to regress out the dominant influence of gDNA contamination and library diversity through linear modeling failed to restore biological clustering (Supplementary fig. 7C). Even after major technical drivers were corrected for, donor phenotypes remained non-separable, while dataset-specific batch effects persisted as the primary organizational feature of the data.

We also assessed other potential confounders such as platelet RNA contamination. Focusing on the gDNA-free datasets, we confirmed that the platelet RNA fraction is a strong contributor to cell-free transcriptome profiles, although its influence was not as large as gDNA contamination or library diversity (Fig 5E). Within datasets, the gene abundance-based PCA also revealed clusters of samples consistent with the groupings observed in the cell type of origin analysis (Supplementary figs. 7D-G).

Finally, we evaluated the influence of microbial sequences. Since gDNA contamination levels were found to be confounded with microbial contamination in some datasets, we again decided to include only gDNA-free datasets in this analysis. We observed no detectable influence (Fig. 5F), suggesting that microbial reads do not cross-map to the human genome to skew human gene abundance profiles. To verify this, we performed a benchmark by re-quantifying all samples after computationally removing every read classified as non-human by Kraken2. We then compared these ‘human-only’ gene abundances against the original ‘unfiltered’ abundances to determine if the presence of foreign reads skewed the results. We found that the correlation between these ‘human-only’ and ‘unfiltered’ gene abundances is both high (median r=0.96 across all samples, Supplementary fig. 8A), and largely independent of the non-human read fraction (Supplementary fig. 8B), confirming that microbial sequences do not skew human gene quantification. However, this stability is lost in samples with an average effective fragment length (EFL, defined in Supplementary fig. 1B) below 100 bp (median r=0.61), compared to longer fragments (median r=0.96; p < 2.2e-16, Wilcoxon rank sum test, Supplementary fig. 8A). This instability is likely a technical artifact caused by taxonomic mis-assignment: as noted above, shorter fragment lengths significantly impair Kraken2’s classification accuracy, often leading to high fractions of unclassified or misclassified sequences (Supplementary fig. 2). In these cases, genuine human fragments are mistakenly removed during the filtering process, whereas longer fragments (EFL ≥ 100 bp) provide the necessary specificity to ensure robust and accurate phylogenetic assignments by Kraken2.

Remarkably, the variation observed even within the gDNA-free cfRNA-Seq datasets was larger than that observed across 29 different tissues sampled from 4 individuals in ENCODE samples (Fig. 5A). This observation corroborates our hypothesis that the high variability across the datasets studied is mainly due to technical effects.

To provide a more quantitative assessment of these drivers, we performed a variance partition analysis (VPA)^40^ across the entire cohort (Fig. 5G). This analysis confirmed that the Broad Protocol Category (BPC) is the primary determinant of transcriptomic variation, explaining the largest fraction of variance across genes. Significant contributions were also attributed to library diversity (NG80), gDNA contamination levels (FSR) and dataset-specific batch effects, while the donor phenotype accounted for only a negligible fraction of the total variance. Interestingly, we observed that the number of mapped fragments (sequencing depth) retained a substantial influence on variance, indicating that TPM normalization is insufficient to fully eliminate depth-dependent technical noise.

Finally, we investigated whether restricting the analysis to high-quality data could restore biological clustering. We performed a principal component analysis restricted exclusively to gDNA-free plasma samples (N=1,586). Even in this filtered subset, technical factors remained the dominant drivers: the first principal component (PC1) accounted for 60.49% of the total variance and primarily separated samples by protocol and laboratory (Supplementary fig. 7H). When colored by clinical status, samples of different phenotypes remained largely intermingled within their respective technical clusters, and a targeted sub-analysis focusing only on healthy controls (N=979) confirmed that protocol-driven separation persists regardless of donor health. These results demonstrate that removing contaminated samples is insufficient to overcome the lack of standardization in the cfRNA-Seq field.

### Recommendations for cfRNA-Seq studies

Based on our analysis, we provide the following recommendations to guide experimental design, protocol selection and accurate reporting of results in cfRNA-Seq research. We do not cover topics that have already been extensively evaluated by the exRNAQC consortium^11^, such as blood collection tubes,RNA extraction kits, or small RNA profiling.

Our finding that pre-analytical variables and batch effects are the predominant drivers of variation in cfRNA profiles underscores the critical need for homogeneous sample processing within any given study. When uniform processing is not feasible, experimental design must incorporate randomization of samples across variables of interest to avoid confounding results. To foster maximum interpretability and reproducibility, we strongly advise that all experimental variables be exhaustively reported according to available standards, as proposed previously^11,29^. Because our analysis identified gDNA contamination as a major source of technical noise, comprehensive reporting of its proxies, such as the fraction of spliced reads (FSR), is essential.

To help choose the most suitable library preparation protocol, we have compiled our main quality metrics into a comprehensive figure that scores methodologies according to their performance (Fig. 6 and Supplementary fig. 9). This data provides clear, evidence-based guidance about which approach to use under different scenarios. We recommend exome capture based (EB) workflows for studies focusing on known protein-coding genes, as they consistently produced the most diverse and gDNA-free libraries. As noted above, the low gDNA content of EB libraries in this study reflects the DNase digestion step used upstream in the specific workflow shared by all three EB datasets, rather than the exome capture itself. Researchers adopting an EB workflow should not assume equivalent protection against gDNA, unless a comparable digestion step is included in their protocol. Rigorous DNase treatment, not the choice of enrichment chemistry *per se*, is the determinant of a gDNA-free library. On the other hand, EB protocols are ill-suited to more exploratory analyses such as the investigation of microbial signatures, or biomarker discovery outside well-annotated human genes. For such purposes, whole-transcriptome random-primed (WRR) protocols are more suitable. To correct for typical low library diversity of WRR methods, affinity-based depletion of abundant, presumably uninformative genes such as rRNA and srpRNAs, as employed in other studies^1,5,41^, is required. Given the susceptibility of WRR methods to gDNA contamination, rigorous DNase treatment should be considered essential. Our analysis suggests double DNase treatment as a highly effective approach producing the most consistent high-quality libraries among all non-EB protocols. The *in-silico* removal of unspliced reads, a strategy previously proposed to mitigate gDNA contamination^2,5,24^, is often impractical. This approach not only leads to a considerable loss of sequence data, but also systematically discards all genuine single-exon genes from downstream analyses.

**Figure 6.**
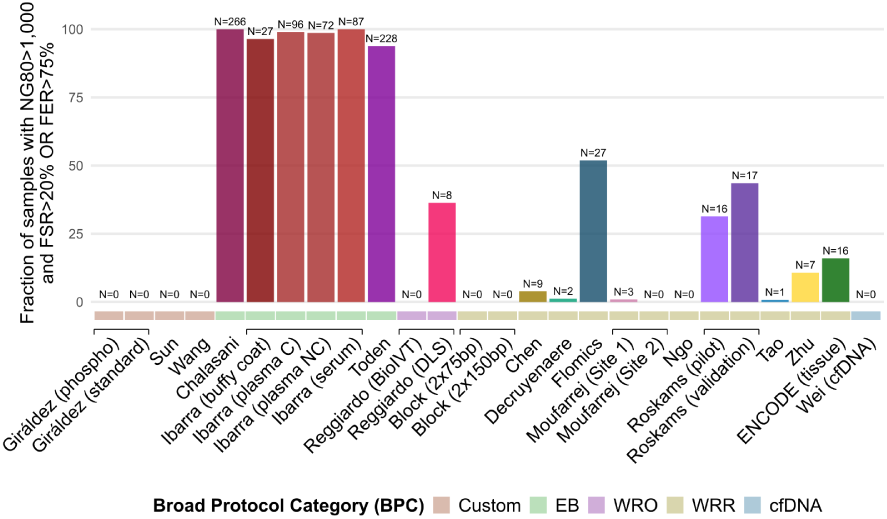
Fraction of high-quality libraries in each dataset. High-quality libraries are defined as those exhibiting an NG80 of more than 1,000 and an FSR greater than 20% or an FER of more than 75%. A more detailed sample survival analysis is available in Supplementary fig. 9.

Conversely, we hesitate to recommend oligo-dT-based (WRO) methods, as they are ill-suited for the degraded nature of cfRNA, and the single representative dataset in our study may have been affected by cellular RNA contamination. We cannot recommend custom protocols at this time, as they often produced low-quality libraries or required idiosyncratic bioinformatics pipeline settings, which hinders cross-study comparison.

Finally, while we find that human gene quantification is generally robust against microbial sequence interference regardless of fragment length, we identify an effective sequencing fragment length (EFL) of at least 100 bp as a requirement for cfRNA-Seq studies involving phylogenetic profiling. Because shorter fragments are highly susceptible to taxonomic mis-assignment (where *e.g.* genuine human reads are either unclassified or misclassified as non-human), longer fragments are essential to ensure the correct call of microbial signatures and to avoid the artifactual removal of human signal during taxonomic filtering.

## Discussion

In this extensive cross-study analysis of 2,356 sequencing libraries, we sought to identify the primary sources of unwanted transcriptomic variation in cfRNA-Seq datasets to help standardize workflows and guide future cfRNA biomarker discovery efforts. Our analysis reveals that the cell-free transcriptome landscape is overwhelmingly shaped not by biological signals, but by technical pre-analytical variables tied to the chosen experimental workflow. This profound heterogeneity, which we show can create more variation than that observed across 29 distinct human tissues, represents a critical challenge to the reproducibility and clinical application of cfRNA-based diagnostics. While other studies have highlighted the importance of technical factors^1,11,12^, ours is the first to systematically investigate this issue at such a comprehensive scale using a controlled, uniform bioinformatics pipeline, leading to several novel findings. Because this uniform analytical approach is optimized for mRNA and lncRNA quantification, our findings and recommendations apply primarily to the long RNA fraction of the cell-free transcriptome.

We demonstrate that the low concentration and highly degraded nature of cfRNA makes it particularly susceptible to contamination from both human and microbial gDNA. While both are prevalent, their impact on data interpretation differs dramatically. We show for the first time on a large scale that human gDNA contamination of cfRNA creates strong biases in human gene expression quantification. In contrast, while microbial sequences are common, they likely have a negligible impact on human gene abundance profiles. Nevertheless, this susceptibility to microbial contamination can confound studies that aim to link microbial signatures to specific phenotypes, and thus requires special attention.

To assess the actual biomarker information content of cfRNA samples, we propose NG80 as a straightforward and intuitive metric for evaluating library diversity. Using this measure, our analysis reveals that cfRNA libraries are prone to being overwhelmed by a few, likely uninformative RNA species. In addition, we confirm the significant influence of platelet RNA contamination on cell-free transcriptome profiles.

Encouragingly, the strong biases introduced by most technical and pre-analytical variables are largely controllable. Their influence can be minimized both prospectively, through the use of optimal, consistent experimental conditions, and retrospectively, by applying computational data quality filters (*e.g.* by eliminating gDNA-contaminated samples, or ignoring lowly-expressed genes), though to a much lower extent.

While our current study focuses on these filters to expose raw technical heterogeneity, we found that even regressing out primary technical drivers (NG80 and FSR) was insufficient to recover a cohesive biological signal. This suggests that the technical noise in cfRNA is deeply embedded across multiple levels. Consequently, while the systematic evaluation of more advanced batch-correction methods remains a vital future perspective, these results suggest that experimental standardization is the only reliable path to robust cfRNA biomarker discovery.

Implementing such standardization in clinical practice, however, remains a primary challenge. Logistical constraints often make it difficult to homogenize procedures between healthy and disease cohorts, which can perpetuate the confounding variables identified in our analysis. By quantifying the impact of these variables, addressing concerning batch effects that can confound results, and providing clear recommendations, this work offers a framework to design more robust, reproducible, and clinically meaningful cfRNA-Seq studies.

## Methods

### Initial processing of external sequencing data

All raw FASTQ files from publicly available datasets were downloaded from the NCBI Sequence Read Archive (SRA). Access to protected datasets (Decruyenaere) was granted upon request by corresponding authors; the corresponding FASTQ files were downloaded from the European Genome-phenome Archive (EGA).

#### Collection and curation of clinical and experimental metadata

Metadata was collected from the corresponding published articles, SRA run metadata, the EGA archive metadata, and GEO series matrix files. The collection center information for the Toden dataset was provided by the authors via the GitHub repository associated with the Vorperian *et al.* study^39^. Phenotype information could not be inferred for a total of 66 samples from the Chalasani, Ibarra and Toden datasets. Both Giráldez batches (“standard” and “phospho-RNA-Seq”) were classified as custom BPC to simplify the analysis. The metadata table was harmonized by aligning column names and standardizing phenotype descriptions (Supplementary table 1).

#### Dataset and sample inclusion criteria

We systematically included cfRNA-Seq datasets published prior to February 2024 that we identified as having publicly accessible raw FASTQ files. The 15 selected studies comprised a total of 2,747 samples after replicate merging, plus 52 internal Flomics samples, bringing the total to 2,799. We excluded those samples not relevant to our analysis for the following reasons:

- Chen *et al.*: *E. coli* and brain samples (n = 3)
- Block *et al.*: Tissue, and plasma-derived vesicles (n = 12).
- Tao *et al.*: Tissue and PBMC, and non-cfRNA-seq assays (MeDIP-Seq and miRNA-Seq) (n = 277).
- Wei *et al.*: Tissue samples (n = 6).
- Wang *et al*.: Only samples from the technology optimization study were publicly available. We excluded samples processed with the adapter ligation-based method NEBNext Small RNA Library Prep Set (n = 3), samples for which modifications were applied to the protocol, such as removing the ExoI enzyme or changing the concentration of the RT primers, as these conditions were not optimal (n = 24).
- Giraldez *et al.*: Synthetic samples (n = 2), protocol optimization samples (n = 30).
- Sun *et al.*: cfDNA samples (n = 4), protocol optimization samples (n = 11).
- Decruyenaere *et al.*: FFPE samples (n = 69).
- Reggiardo *et al.*: Samples sequenced on the ONT platform (n = 2).
- ENCODE: From the EN-TEx resource, non-human, total RNA-seq paired-ended samples were excluded (n = 1,494).

#### Replicate merging

Chalasani and Toden data included technical replicates of the same sample. Those sequencing runs were merged into samples using the information available in SRA and the corresponding paper. Specifically, the “Isolate” column from the provided SRA metadata was used to identify replicates and merge their FASTQs into a single sample-specific FASTQ.

### Confounding of experimental variables with phenotype

We calculated the association between sample phenotypes and technical variables across datasets using Cramér’s V. For each dataset, we computed pairwise Cramér’s V between a simplified phenotype label (healthy, cancer or non-cancer disease) and 11 technical factors (e.g., collection center, centrifugation steps, read length). One sample from the Toden dataset with an unknown collection center was removed to allow computation.

### Flomics cfRNA-Seq library preparation

Eight plasma samples from the Flomics dataset were provided by the Hospital Clínic de Barcelona - IDIBAPS Biobank (HCB-IDIBAPS, B.0000575, Platform ISCIII Biobanks and Biomodels), Spain. 44 samples were provided by the Andalusian Public Health System Biobank (BBSSPA). All samples were processed following standard operating procedures with the appropriate approval of the Ethics and Scientific Committees. In accordance with the international ethical and legal framework, the release of biological samples and associated data from the HCB-IDIBAPS biobank is approved by both the internal and external Scientific Committees, as well as by the Ethics Committee from HCB. The IDIBAPS release was granted approval code A3-C22016. BBSSPA sample release was approved by the *Portal de Ética de la Investigación Biomédica de Andalucía (PEIBA)* with codes 2042-N-22 and SICEIA-2024-002798.

In both cases, whole blood was collected in EDTA collection tubes and processed to plasma within four hours. Whole blood was centrifuged at 1,500 x g for 10 minutes at room temperature to separate plasma. The plasma layer was transferred to a separate tube and centrifuged at 2,500 x g for 15 minutes at room temperature to minimize platelet content. 1 mL aliquots of the double centrifuged plasma were stored at −80 °C. Prior to cfRNA isolation plasma aliquots were thawed and centrifuged at 400 x g for 2 minutes at 4 °C to remove precipitate. 1.5 pg of External RNA Controls Consortium (ERCC) RNA Spike-In Mix (ThermoFisher) was added to each 1 ml of plasma. cfRNA was isolated from 1 mL of plasma using the Maxwell miRNA Plasma and Serum Isolation kit (Promega). Genomic DNA was removed using bovine DNase I (Promega) and HL-dsDNase (ArcticZymes Technologies). Library preparation was performed using the SMARTer Stranded Total RNA-Seq Kit v3 - Pico Input Mammalian (Takara Bio). Libraries were sequenced on the NextSeq 2000 platform (Illumina) with a read length of 2×150bp.

### Primary and secondary bioinformatics analysis

All RNA-Seq datasets were processed using the nf-core/rnaseq^31^ Nextflow^32^ pipeline (version 3.8.1, Fig. 1D) from raw data (FASTQ files), using STAR^33^ as the read mapping software and Salmon^34^ to quantify annotated genes. All samples were processed using the same custom STAR parameters (“--readFilesCommand zcat --twopassMode None --outSAMprimaryFlag OneBestScore --outFilterMismatchNmax 999 --outFilterMismatchNoverReadLmax 0.04 --alignIntronMin 20 --alignIntronMax 1000000 --alignMatesGapMax 1000000 –outFilterType BySJout --alignSJDBoverhangMin 3 --alignSJoverhangMin 8 --peOverlapNbasesMin 40 --peOverlapMMp 0.8 --quantMode TranscriptomeSAM --quantTranscriptomeBan Singleend --outSAMattributes All”). Pipeline parameters were adapted to the cDNA library preparation method used in each dataset where appropriate (Supplementary table 2). Nextflow configuration files are available in the project’s GitHub repository (https://github.com/Flomics/fl-cfRNAmeta), under the nextflow/ subdirectory. We used genome assembly version GRCh38 coupled with the GENCODE v39 annotation^42^. We incorporated the synthetic SIRV-set 3 spike-in transcripts (including the ERCC mix) to both the reference genome and annotation.

Mapped fragments were deduplicated using samtools “markDuplicates” or umiTools “UMI-dedup”, depending on the presence of UMIs in the library construction methods. The fraction of spliced reads (FSR) was calculated as the fraction of uniquely mapped, deduplicated reads that span introns (i.e., with an “N” in their CIGAR string). The fraction of exonic reads (FER) was calculated as the fraction of STAR-mapped and deduplicated reads with at least one base of overlap with a GENCODE-annotated exon (excluding synthetic spike-in sequences). The sequencing data strandedness (namely, the fraction of fragments mapping to the correct gene orientation) was calculated using the “inferExperiment” module from the “RSeQC” package^43^.

Kraken2^35^ (version 2.1.2) was run separately from the nf-core/rnaseq pipeline to infer the taxonomic composition of samples using fl-kraken2, a Nextflow pipeline developed in house (https://github.com/Flomics/fl-kraken2). FASTQs were matched against the pre-computed PlusPF sequence index (version 2023-03-14, downloaded from https://benlangmead.github.io/aws-indexes/k2, k-mer size: 35). The PlusPF includes the following sequence databases: RefSeq archaea, bacteria, viral, plasmid, human, UniVec_Core, protozoa and fungi. All Kraken2 parameters were set to default. To study the influence of non-human reads on gene abundance profiles, we used fl-KrakenBAM (https://github.com/Flomics/fl-KrakenBAM), a Nextflow pipeline developed in house to extract human reads from BAM files based on Kraken2 classifications. fl-KrakenBAM generates Salmon-compatible transcriptome BAM files devoid of non-human reads according to Kraken2 classification.

### Biotype representation

Based on Ensembl/GENCODE gene-biotype annotation^42^, a custom biotype classification was created to highlight relevant categories in the context of cfRNA. Labels for specific Ensembl biotypes were merged into more general categories: protein-coding transcripts were relabeled as “mRNA”, various pseudogenes subtypes combined under the category “Pseudogene”, related small RNAs such as snRNA and snoRNA were grouped together, whereas other small RNAs types (scaRNA, sRNA, scRNA, vault RNA and MT tRNA) were classified into “Other small RNAs”. Finally, signal recognition particle RNAs (srpRNA, a subtype of misc_RNA consisting in genes RN7SL1, RN7SL2, and RN7SL3), mitochondrial ribosomal RNAs (MT rRNA) and nuclear ribosomal RNA (rRNA) were categorized separately. ERCC spike-ins were excluded from this analysis. A detailed mapping between the Ensembl biotypes and our custom biotype categories is provided in Supplementary table 3.

Gene TPM (transcripts per million) counts were produced by nf-core/rnaseq using Salmon^34^ (see above). The total aggregated expression per biotype was calculated for each sample by summing the TPM values of all genes belonging to each biotype category. For visualization and comparative analysis, aggregated gene expression values per biotype were converted to percentages, representing the proportion of total TPM per sample attributed to each biotype. Mean values per biotypes across multiple samples in each experimental batch were calculated to facilitate comparisons.

### Taxonomic distribution analysis

The detailed reports generated by Kraken2 for each sample were collected. The NCBI taxon “other sequences”, which contains spike-in and synthetic sequences (vector, plasmid, etc), was excluded from the analysis. For each sample, read counts statistics were aggregated into a simplified list of mutually exclusive categories: human (Homo sapiens), fungi (Fungi), other_eukaryotes (Eukaryota except human and Fungi), bacteria (Bacteria), other (Archaea, Viruses, and unspecific taxa above the domain rank), and unclassified.

### Microbial reads cross-mapping analysis

We used fl-kraken2, an in-house developed Nextflow pipeline for the taxonomic profiling of whole FASTQ files (URL: https://github.com/Flomics/fl-kraken2). fl-KrakenBAM (URL: https://github.com/Flomics/fl-KrakenBAM) was used to filter non-human reads from transcriptome BAM files. These “human-only” BAM files were subsequently used to re-quantify genes using nf-core/rnaseq using the same parameters as for the “unfiltered” BAM files obtained previously.

Gene count matrices were log-transformed (log10(raw_count + 1)) after excluding ERCC and SIRV spike-in records. For each biological sample, we calculated the Pearson correlation coefficient (r) between “human-only” and “unfiltered” reads-based quantifications. To statistically quantify the impact of effective fragment length (EFL) on quantification consistency, samples were dichotomized using a 100bp average EFL threshold, and differences in the distribution of r values between groups were evaluated using a Wilcoxon rank sum test. The full analysis script is available in the project’s GitHub repository (https://github.com/Flomics/fl-cfRNAmeta/blob/dev/src/correlation_analysis.R).

### RNA library diversity analysis

The NG80 metric (number of genes accounting for 80% of a sequencing library’s mapped and deduplicated reads) was calculated using the Salmon raw counts matrix as input, as produced by the standard nf-core/rnaseq pipeline. The full script for the calculation of NG80 can be found on the code repository (https://github.com/Flomics/fl-cfRNAmeta/blob/dev/src/ng80.R). NP80 (number of genes accounting for 80% of protein-coding reads) values were calculated using the same procedure, after pre-filtering the raw counts matrix to keep only those genes that are classified as “mRNA”, according to the custom biotype classification mentioned above.

Additional diversity metrics were computed on the same count matrices as NG80, using R libraries “vegan” version 2.7-2 (Shannon entropy) and “ineq” version 0.2-13 (Gini index) (see also diversity_scatterplots.R wrapper script in the provided github repository).

### Cell type of origin deconvolution

The nu-SVR deconvolution method, as developed by Vorperian *et al.*^39^, was employed to predict the relative cell type contribution in cfRNA. We used Tabula Sapiens^44^ 1.0, as provided in the Vorperian *et al.* paper, as the cell type basis matrix for nu-SVR. Only gDNA-free cfRNA-Seq datasets were analyzed. We defined “major cell types” (MCT) on a per-dataset basis as cell types with a relative abundance of at least 10% in at least 10% of the samples. Subsequently, a global list was generated by taking the union of all major cell types identified across the individual datasets. This global MCT list consisted of platelets, monocytes, erythrocytes, B cells, macrophages and neutrophils, and was kept for further analysis.

### Dimensionality reduction

Dimensionality reduction of gene expression data was performed using Principal Component Analysis (PCA) on Salmon-based TPM values. Spike-in control sequences were explicitly excluded from this analysis to prevent additional technical variance from biasing the results. To stabilize variance and reduce the skewness of the data, TPM values were log-transformed (log(TPM +1)) prior to PCA. PCA was implemented using the scikit-learn Python library. The first two principal components were used to embed samples in a two-dimensional space, visually representing sample similarity and coloring them based on different metadata properties or QC metrics.

A similar analysis was performed using t-distributed Stochastic Neighbor Embedding (tSNE) with identical data pre-processing. Various perplexity parameters were evaluated (20, 50, 100), and a value of 20 was chosen for Supplementary fig. 7A, given that all led to similar results.

### Variance partition analysis

Variance partition analysis was performed using the “variancePartition” R package version 1.40.2 to quantify the contribution of technical and biological variables to gene expression variance across samples. The full analysis script is available in the project’s GitHub repository (src/variance_partition.R).

Gene expression data was obtained from the pre-computed TPM matrix. Genes with TPM > 1 in fewer than 10% of samples were excluded prior to modelling, retaining 18,985 genes. Expression values were log_2_(TPM + 1) transformed before fitting. The analysis was done on the full cohort of 2,356 samples.

Candidate variables for the model were all pre-analytical variables reported in Fig 1A, plus analytical variables identified as relevant: NG80, FER, FSR, platelet contribution, and mapped reads (“Mapped fragments”). Collinearity between these variables was assessed using the “canConPairs” function. Variables with an RMS canonical correlation higher than 0.999 with either BPC or Dataset, as detected by the function, were excluded from the model.

The remaining variables were modelled as follows: continuous variables (NG80, FER, FSR, platelet contribution, and mapped reads) were included as fixed effects. Categorical variables (Dataset, Phenotype, and BPC) were modelled as random effects. The model was fitted independently for each gene using “fitExtractVarPartModel”, and the results plotted using ggplot’s “geom_violin” function.

### Batch effect correction by linear modeling

Gene expression values were corrected for major sources of unwanted technical variation using a linear-model to adjust for NG80 and FSR. Only plasma cfRNA-Seq datasets were taken into account in this analysis (*i.e.*, excluding “Ibarra buffy coat”, “Ibarra serum” as well as “Wei” (cfDNA) and ENCODE (tissue RNA). Filtered count matrices were normalized by the trimmed mean of M-values method and transformed to log2 counts per million using normalized library sizes. For each gene, a linear model was fitted with limma, modeling log2-TMM expression as a function of disease status, which was retained as the biological variable of interest, and the technical covariates selected for correction. In the final correction model, the main unwanted sources of variation were NG80 and FSR. Sex was also included when variable across samples. Coefficients corresponding to the unwanted covariates were extracted while excluding the intercept and disease-status terms, and the expression component explained by NG80 and FSR was estimated for each gene and sample from the corresponding design matrix. Corrected expression values were then obtained by subtracting this unwanted-variation component from the original log2-TMM expression matrix, yielding expression values that preserve disease-status-associated signal while reducing variation attributable to library diversity and gDNA contamination technical effects. The effectiveness of the correction was evaluated by principal component analysis and by assessing correlations between the leading principal components and technical covariates before and after correction.

## Data availability

Sequencing data for the Flomics dataset were deposited in the European Nucleotide Archive (ENA) under accession PRJEB90290. Only the samples identified as “flomics_2” in the “sample_alias” field were used in this analysis. All external sequencing datasets used in this paper are available from public repositories, as specified in the corresponding original study.

## Code availability

All code produced as part of this analysis is located in GitHub at the following URLs: https://github.com/Flomics/fl-cfRNAmeta (analysis code), https://github.com/Flomics/fl-kraken2 (Kraken2-based taxonomic profiling) and https://github.com/Flomics/fl-KrakenBAM (extraction of human reads from BAM files based on Kraken2 classification).

## Supplementary materials

### Supplementary figures

Provided as a separate document, with legends.

### Supplementary tables

Supplementary table 1

Curated sample metadata information

Supplementary table 2

RNA-Seq pipeline custom parameters, with URL pointers to Nextflow configuration files

Supplementary table 3

Mapping of ENSEMBL/GENCODE biotype terms

## Acknowledgements

We thank paper authors we contacted for their cooperation and fruitful discussions. We are indebted to the HCB-IDIBAPS Biobank for scientific advice and sample and data procurement. The project ‘LiquiDx’ (reference CPP2021-008897) is financed by the Spanish Ministry of Science, Innovation and Universities (MCIN/AEI/10.13039/501100011033) and by the European Union “NextGenerationEU/PRTR”. We thank the partners of the LiquiDx project for scientific advice and sample and data procurement: Health and Biomedicine, Leitat Technological Center, Barcelona Science Park, Barcelona 08028, Spain and the Andalusian Public Health System Biobank, Coordinating Node, 18016 Granada, Spain. We are grateful to Sílvia Pérez-Lluch (Centre for Genomic Regulation (CRG), Barcelona) for her scientific advice. CT and GA are supported by the Industrial Doctorates Plan of the Department of Research and Universities of the Government of Catalonia (references 2022-DI-00108 and 2024-DI-00013, respectively). JL, MW, PS and PM are supported by the Torres Quevedo programme (references PAQ20-011298, PAQ2022-012611, PTQ2022-012612 and PTQ2023-012932, respectively), which is financed by the Spanish Ministry of Science, Innovation and Universities (MCIN/AEI/10.13039/501100011033). We acknowledge support of the Spanish Ministry of Science and Innovation through the Centro de Excelencia Severo Ochoa (CEX2020-001049-S, MCIN/AEI/10.13039/501100011033), and the Generalitat de Catalunya through the CERCA programme. We are grateful to the CRG Core Technologies Programme for their support and assistance in this work.

## Author contributions

Conceptualization: JL

Data Curation: CT-D, MW, JL

Formal analysis: CT-D, GA, PM-M, ECR, MW, JL

Funding acquisition: PM-M, PS, MW, JL

Investigation: CT-D, GA, PM-M, ECR, LC, LG, LS, BN, PS, MW, JL

Methodology: JL, MW, PM-M, ECR

Project administration: JL

Resources: PS, JL

Software: CT-D, GA, PM-M, ECR, LC, MW, JL

Supervision: JL

Visualization: CT-D, GA, PM-M, ECR, MW, JL

Writing – original draft: JL

Writing – review & editing: JL, MW, PS

