## Supplementary Figures for "Systematic cross-study assessment of RNA-Seq experimental workflows for plasma cell-free transcriptome profiling"

### Table of contents


Table of contents 1
Supplementary figure 1 2
- A 3
- B 4
- C 4
- D 5

Supplementary figure 2 6
- A 6
- B 7
- C 7
- D 8
- E 9
- F 9
- G 10

Supplementary figure 3 11
Supplementary figure 4 12
- A 12
- B 13

Supplementary figure 5 14
Supplementary figure 6 15
- A 15
- B 15
- C 16
- D 16
- E 16
- F 17
- G 17
- H 17
- I 18

Supplementary figure 7 19
- A 19
- B 20
- C 20

D 21
E 22
F 22
G 23
H 23
I 24
Supplementary figure 8 25
A 26
B 26
Supplementary figure 9 28
A 28
B 29
C 29

### Supplementary figure 1

**Basic sequencing and genome mapping statistics across datasets. (A)** Sample sequencing depth. Number of sequencing fragments per dataset. Each data point represents a sample. **(B)** Effective sequencing fragment length (EFL). Namely, the "Average Mapped Length" statistic generated by the STAR mapper. It reflects the length of sequencing fragments that the aligner was able to align to the human genome, hence effectively subtracting adapter- and quality-trimmed sequences, as well as overlapping sequences between sequencing mates. **(C)** Fraction of exonic reads (FER). Reads mapping to spike-in sequences were excluded from this analysis. **(D)** Data strandedness. Namely, the fraction of exonic reads that map to the correct annotated strand of the underlying gene. It is expected to be close to 100% for stranded RNA libraries, and to tend towards 50% for both unstranded and genomic libraries. Values substantially below 50% (*e.g.* Ngo dataset) might indicate read 1 / read 2 FASTQ mate swapping error in the input data downloaded from SRA.

A

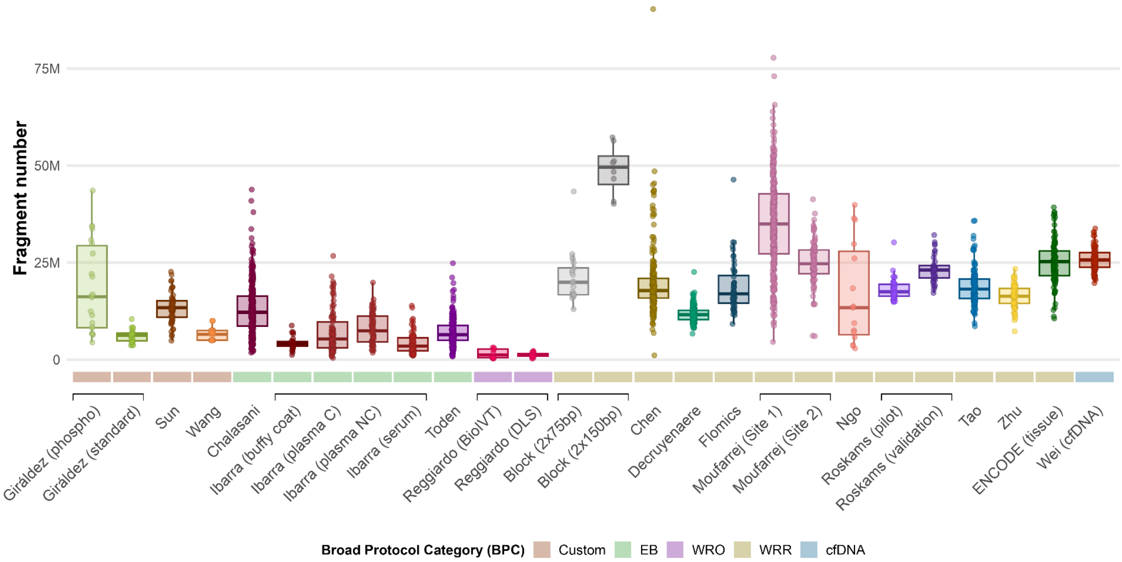

Fragment number
75M
50M
25M
0
Giráldez (phospho)
Giráldez (standard)
Sun
Wang
Chalasani
Ibarra (buffy coat)
Ibarra (plasma C)
Ibarra (plasma NC)
Ibarra (serum)
Toden
Reggiardo (BioIVT)
Reggiardo (DLS)
Block (2x75bp)
Block (2x150bp)
Chen
Decruyenaere
Flomics
Moufarrej (Site 1)
Moufarrej (Site 2)
Ngo
Roskams (pilot)
Roskams (validation)
Tao
Zhu
ENCODE (tissue)
Wei (cfDNA)
Broad Protocol Category (BPC)
Custom
EB
WRO
WRR
cfDNA

B

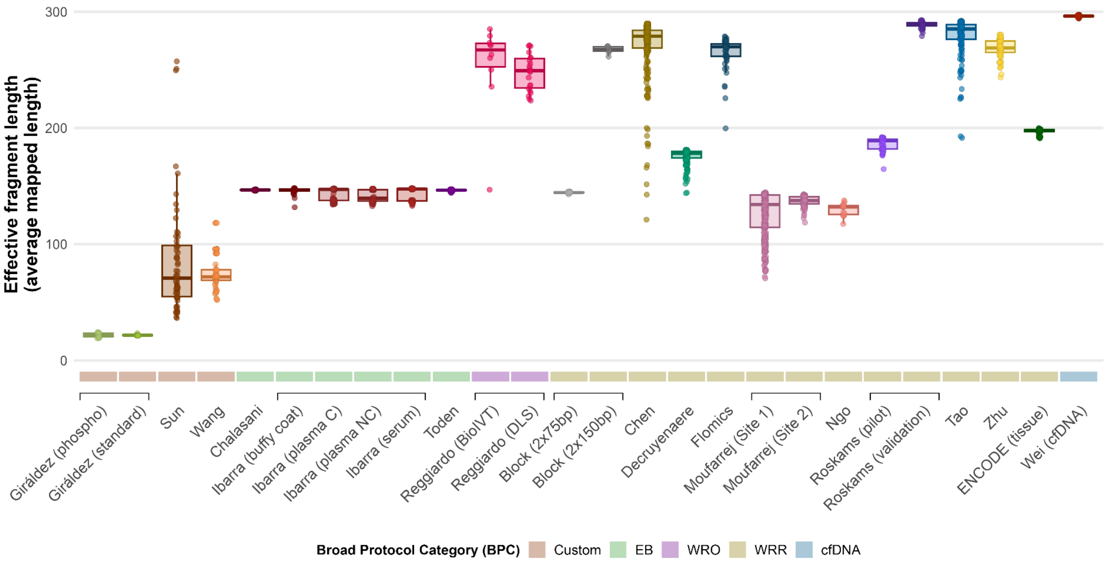
Effective fragment length (average mapped length)
300
200
100
0
Giráldez (phospho)
Giráldez (standard)
Sun
Wang
Chalasani
Ibarra (buffy coat)
Ibarra (plasma C)
Ibarra (plasma NC)
Ibarra (serum)
Toden
Reggiardo (BioIVT)
Reggiardo (DLS)
Block (2x75bp)
Block (2x150bp)
Chen
Decruyenaere
Flomics
Moufarrej (Site 1)
Moufarrej (Site 2)
Ngo
Roskams (pilot)
Roskams (validation)
Tao
Zhu
ENCODE (tissue)
Wei (cfDNA)
Broad Protocol Category (BPC)
Custom
EB
WRO
WRR
cfDNA

C

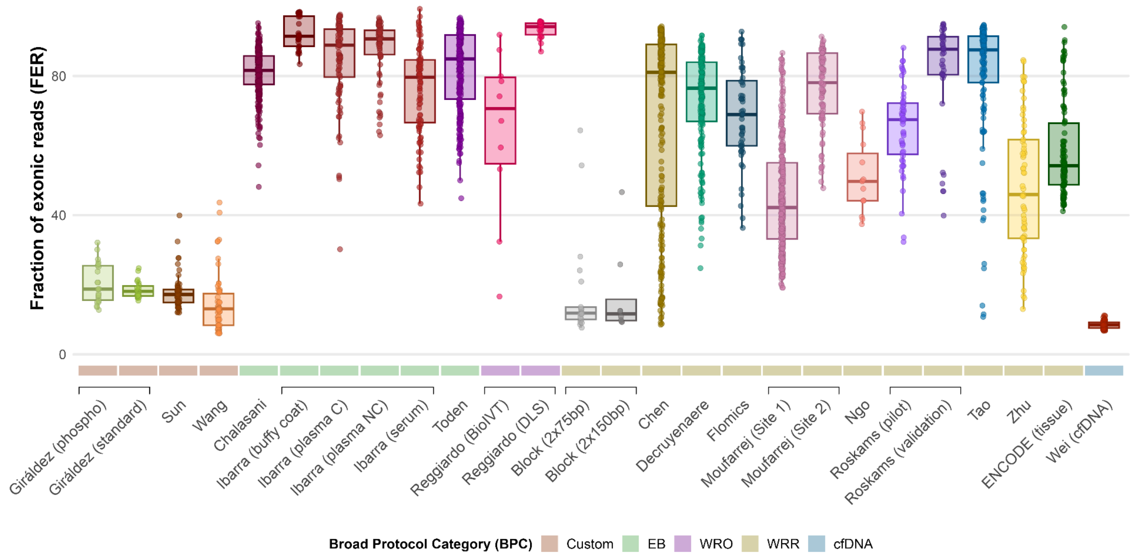
Fraction of exonic reads (FER)
80
40
0
Giráldez (phospho)
Giráldez (standard)
Sun
Wang
Chalasani
Ibarra (buffy coat)
Ibarra (plasma C)
Ibarra (plasma NC)
Ibarra (serum)
Toden
Reggiardo (BioIVT)
Reggiardo (DLS)
Block (2x75bp)
Block (2x150bp)
Chen
Decruyenaere
Flomics
Moufarrej (Site 1)
Moufarrej (Site 2)
Ngo
Roskams (pilot)
Roskams (validation)
Tao
Zhu
ENCODE (tissue)
Wei (cfDNA)
Broad Protocol Category (BPC)
Custom
EB
WRO
WRR
cfDNA

D

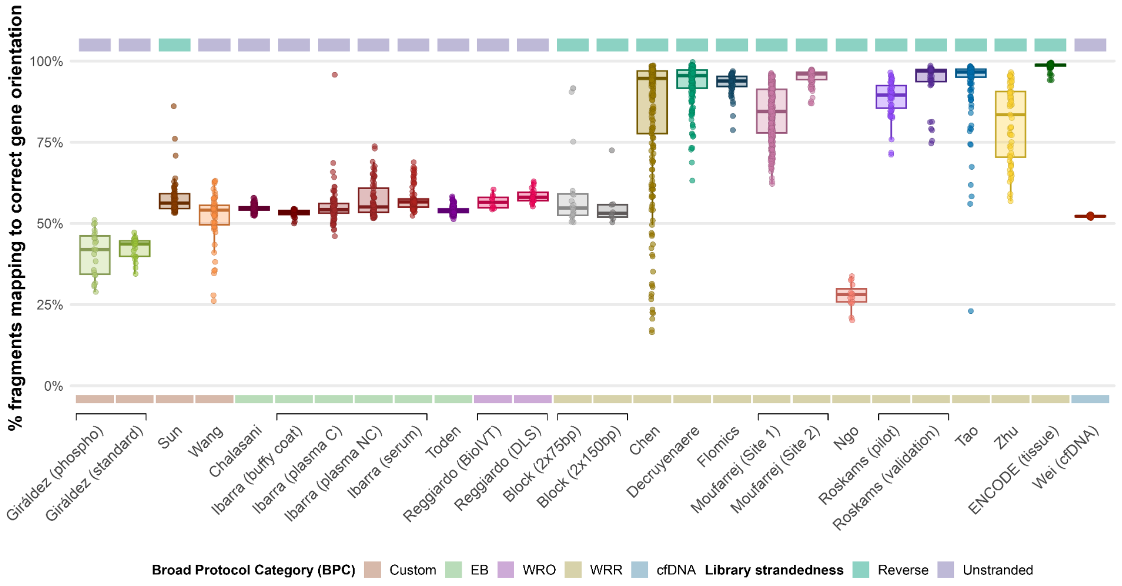

% fragments mapping to correct gene orientation
100%
75%
50%
25%
0%
Giráldez (phospho)
Giráldez (standard)
Sun
Wang
Chalasani
Ibarra (buffy coat)
Ibarra (plasma C)
Ibarra (plasma NC)
Ibarra (serum)
Toden
Reggiardo (BioIVT)
Reggiardo (DLS)
Block (2x75bp)
Block (2x150bp)
Chen
Decruyenaere
Flomics
Moufarrej (Site 1)
Moufarrej (Site 2)
Ngo
Roskams (pilot)
Roskams (validation)
Tao
Zhu
ENCODE (tissue)
Wei (cfDNA)
Broad Protocol Category (BPC)
Custom
EB
WRO
WRR
cfDNA
Library strandedness
Reverse
Unstranded

### Supplementary figure 2

**Variability in taxonomic profiles across samples.** Distribution of the taxonomic profiles of all samples, grouped into simplified categories: **(A)** human (Homo sapiens), **(B)** fungi (Fungi), **(C)** other_eukaryotes (namely, Eukaryota except human and Fungi), **(D)** bacteria (Bacteria), **(E)** other (Archaea, Viruses, and other unspecific taxa), and **(F)** unclassified. Each of the plots shows the distribution of the percentage of reads for one of the categories across samples for each dataset. **(G)** For datasets with available metadata, distribution of bacterial reads across samples grouped by collection center within each dataset.

A

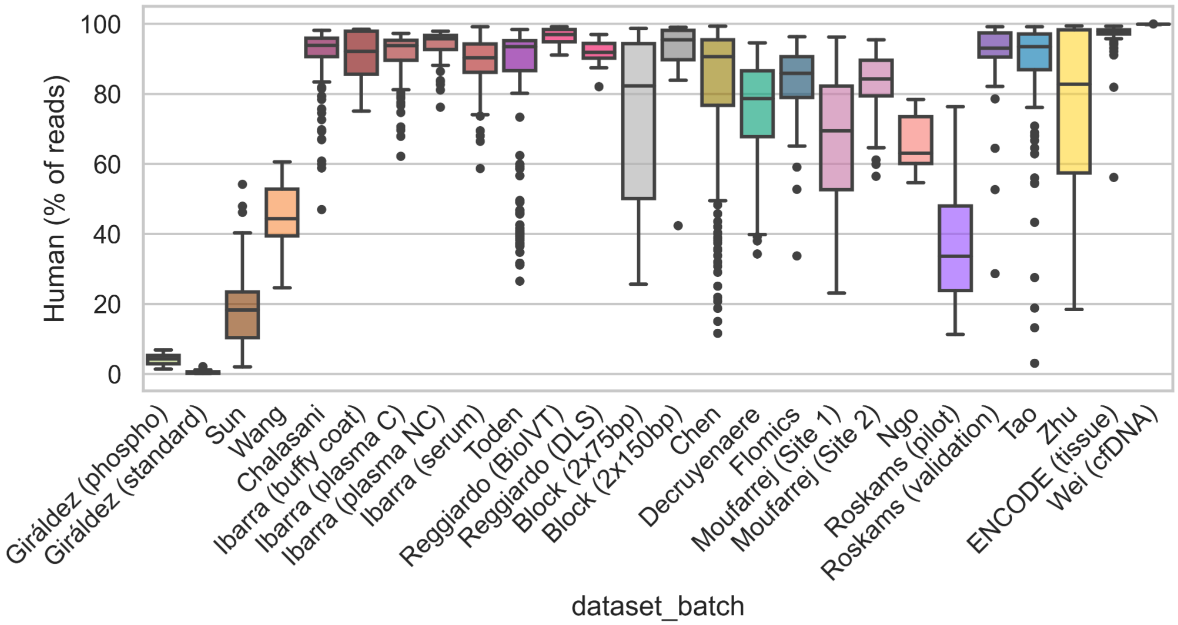

B

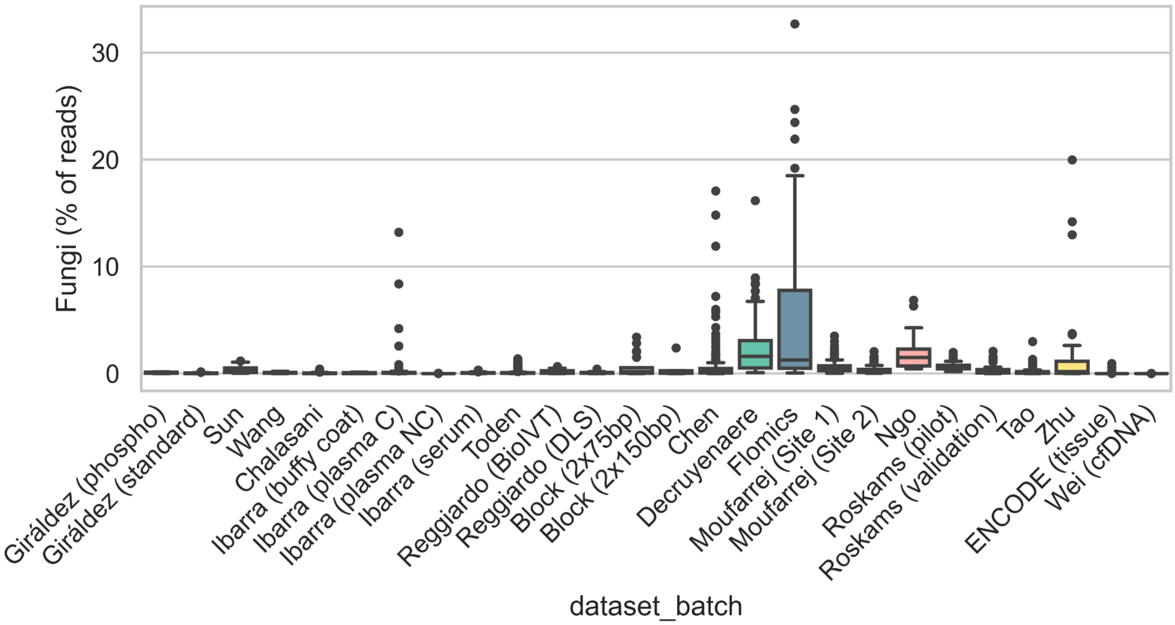

Fungi (% of reads)
30
20
10
0
Giráldez (phospho)
Giráldez (standard)
Sun
Wang
Chalasani
Ibarra (buffy coat)
Ibarra (plasma C)
Ibarra (plasma NC)
Ibarra (serum)
Toden
Reggiardo (BioIVT)
Reggiardo (DLS)
Block (2x75bp)
Block (2x150bp)
Chen
Decruyenaere
Flomics
Moufarrej (Site 1)
Moufarrej (Site 2)
Ngo
Roskams (pilot)
Roskams (validation)
Tao
Zhu
ENCODE (tissue)
Wei (cfDNA)
dataset_batch


C

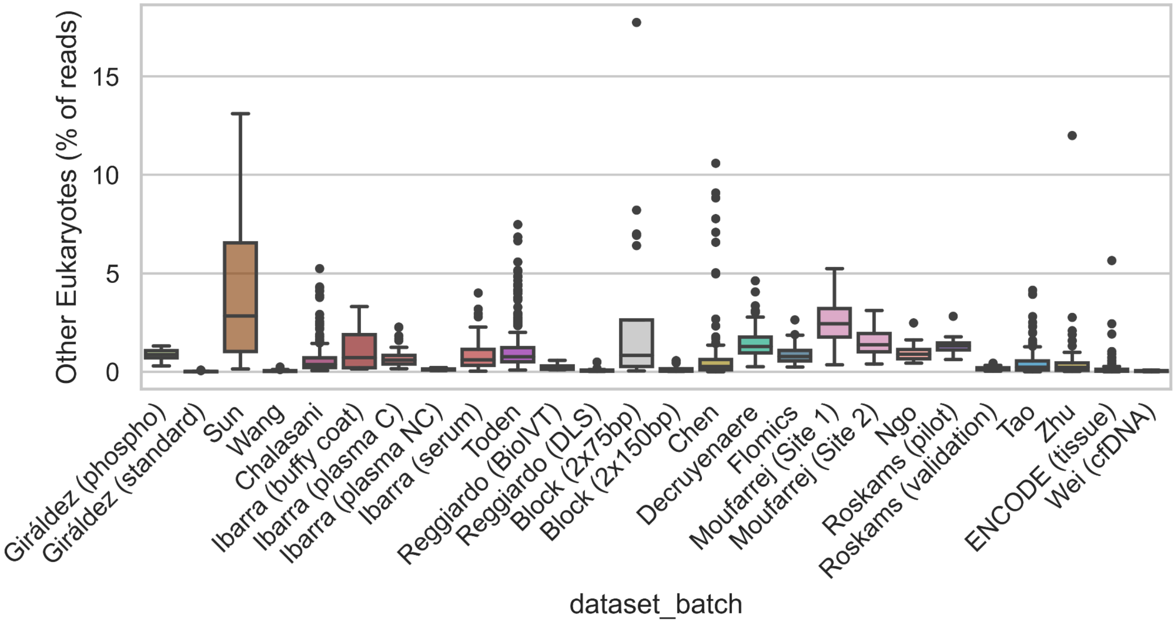

Other Eukaryotes (% of reads)
15
10
5
0
Giráldez (phospho)
Giráldez (standard)
Sun
Wang
Chalasani
Ibarra (buffy coat)
Ibarra (plasma C)
Ibarra (plasma NC)
Ibarra (serum)
Toden
Reggiardo (BioIVT)
Reggiardo (DLS)
Block (2x75bp)
Block (2x150bp)
Chen
Decruyenaere
Flomics
Moufarrej (Site 1)
Moufarrej (Site 2)
Ngo
Roskams (pilot)
Roskams (validation)
Tao
Zhu
ENCODE (tissue)
Wei (cfDNA)
dataset_batch

D

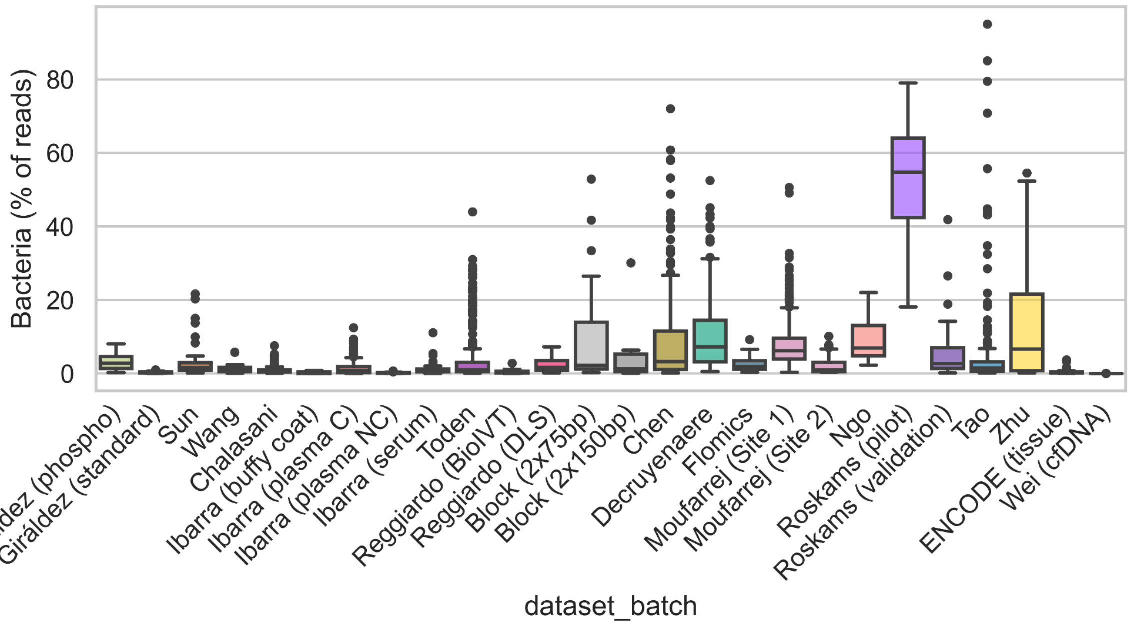
Bacteria (% of reads)
0
20
40
60
80
Giráldez (phospho)
Giráldez (standard)
Sun
Wang
Chalasani
Ibarra (buffy coat)
Ibarra (plasma C)
Ibarra (plasma NC)
Ibarra (serum)
Toden
Reggiardo (BioIVT)
Reggiardo (DLS)
Block (2x75bp)
Block (2x150bp)
Chen
Decruyenaere
Flomics
Moufarrej (Site 1)
Moufarrej (Site 2)
Ngo
Roskams (pilot)
Roskams (validation)
Tao
Zhu
ENCODE (tissue)
Wei (cfDNA)
dataset_batch

E

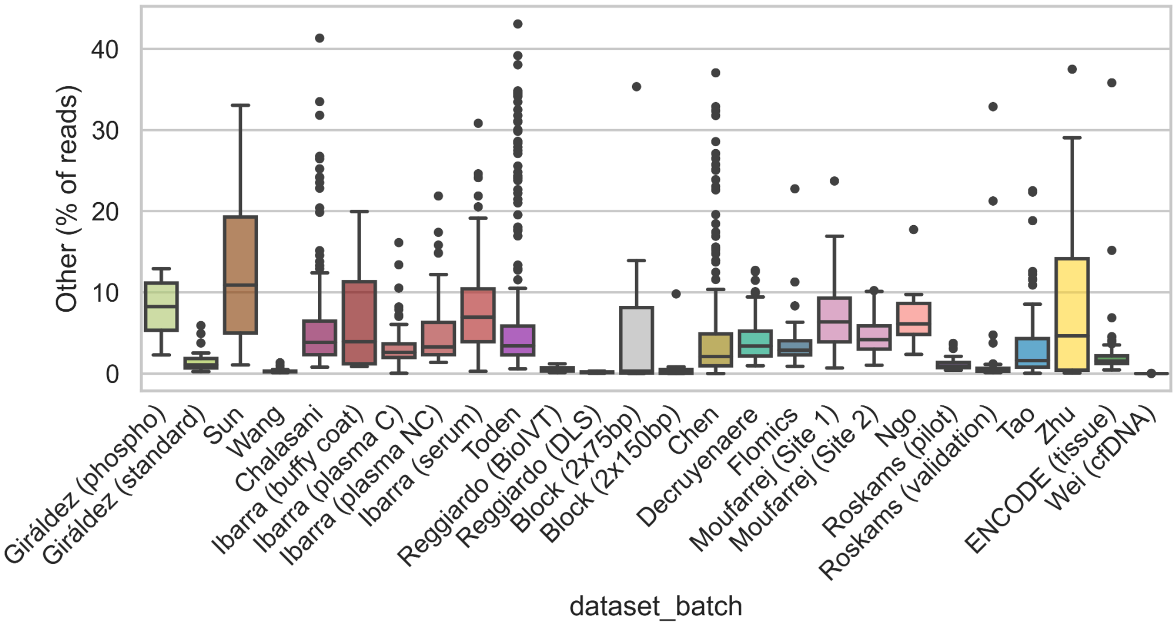
Other (% of reads)
0
10
20
30
40
Giráldez (phospho)
Giráldez (standard)
Sun
Wang
Chalasani
Ibarra (buffy coat)
Ibarra (plasma C)
Ibarra (plasma NC)
Ibarra (serum)
Toden
Reggiardo (BioIVT)
Reggiardo (DLS)
Block (2x75bp)
Block (2x150bp)
Chen
Decruyenaere
Flomics
Moufarrej (Site 1)
Moufarrej (Site 2)
Ngo
Roskams (pilot)
Roskams (validation)
Tao
Zhu
ENCODE (tissue)
Wei (cfDNA)
dataset_batch

F

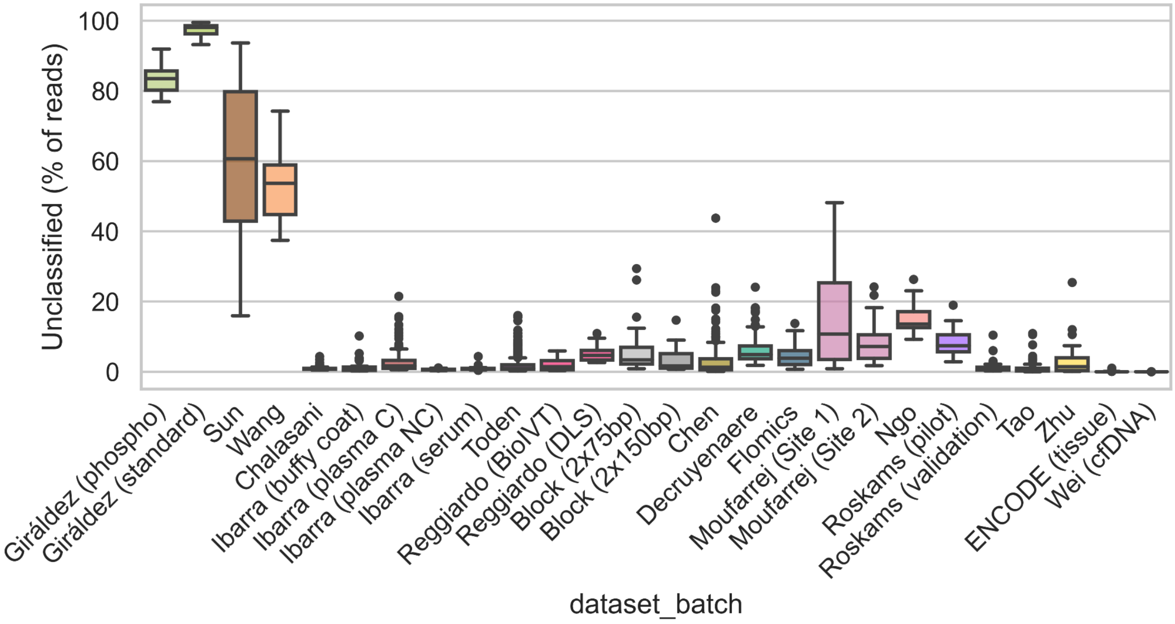
Unclassified (% of reads)
0
20
40
60
80
100
Giráldez (phospho)
Giráldez (standard)
Sun
Wang
Chalasani
Ibarra (buffy coat)
Ibarra (plasma C)
Ibarra (plasma NC)
Ibarra (serum)
Toden
Reggiardo (BioIVT)
Reggiardo (DLS)
Block (2x75bp)
Block (2x150bp)
Chen
Decruyenaere
Flomics
Moufarrej (Site 1)
Moufarrej (Site 2)
Ngo
Roskams (pilot)
Roskams (validation)
Tao
Zhu
ENCODE (tissue)
Wei (cfDNA)
dataset_batch

G

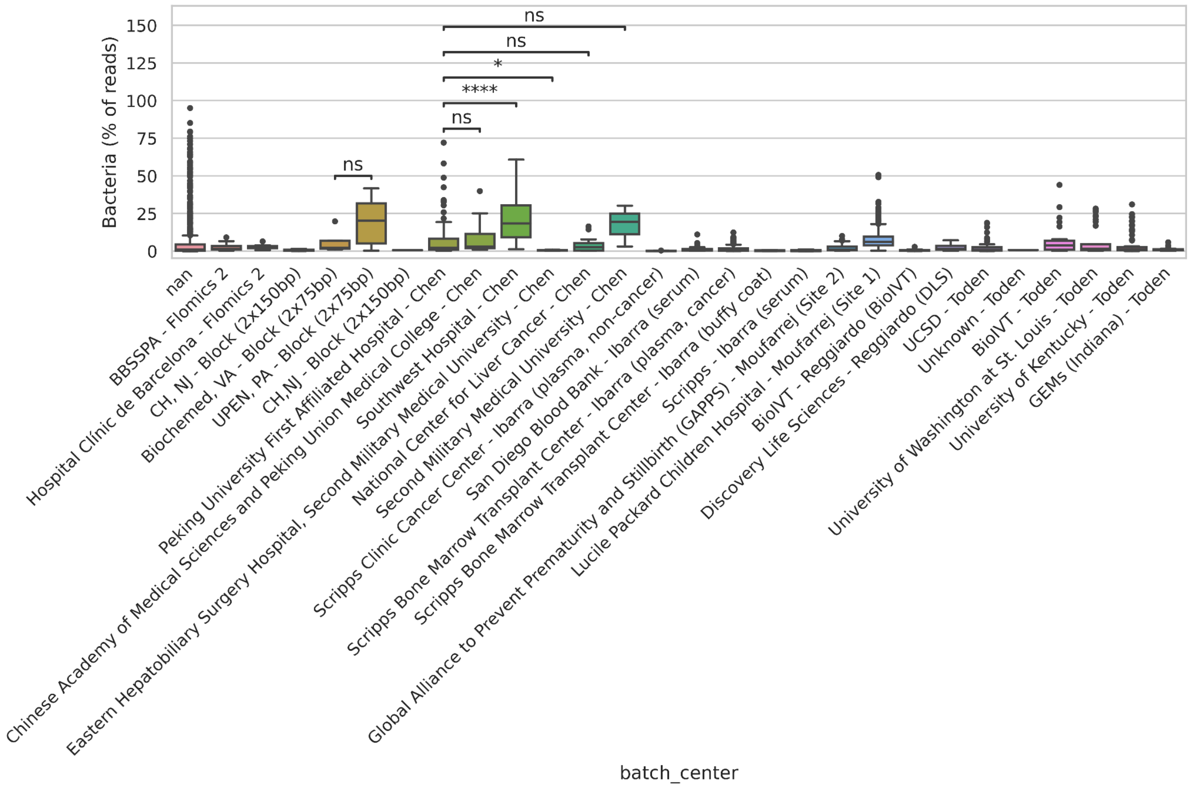

ns
ns
*
****
ns
ns
150
125
100
75
50
25
0
Bacteria (% of reads)
nan
BBSSPA - Flomics 2
Hospital Clínic de Barcelona - Flomics 2
CH, NJ - Block (2x150bp)
Biochemed, VA - Block (2x75bp)
UPEN, PA - Block (2x75bp)
CH,NJ - Block (2x150bp)
Peking University First Affiliated Hospital - Chen
Chinese Academy of Medical Sciences and Peking Union Medical College - Chen
Southwest Hospital - Chen
Eastern Hepatobiliary Surgery Hospital, Second Military Medical University - Chen
National Center for Liver Cancer - Chen
Second Military Medical University - Chen
Scripps Clinic Cancer Center - Ibarra (plasma, non-cancer)
San Diego Blood Bank - Ibarra (serum)
Scripps Bone Marrow Transplant Center - Ibarra (plasma, cancer)
Scripps Bone Marrow Transplant Center - Ibarra (buffy coat)
Scripps - Ibarra (serum)
Global Alliance to Prevent Prematurity and Stillbirth (GAPPS) - Moufarrej (Site 2)
Lucile Packard Children Hospital - Moufarrej (Site 1)
BioIVT - Reggiardo (BioIVT)
Discovery Life Sciences - Reggiardo (DLS)
UCSD - Toden
Unknown - Toden
BioIVT - Toden
University of Washington at St. Louis - Toden
University of Kentucky - Toden
GEMs (Indiana) - Toden
batch_center

### Supplementary figure 3

**Correlation between human genome mapping rate (X axis) and fraction of microbial reads (Y axis) per dataset and batch**. Each dot corresponds to a sequencing library

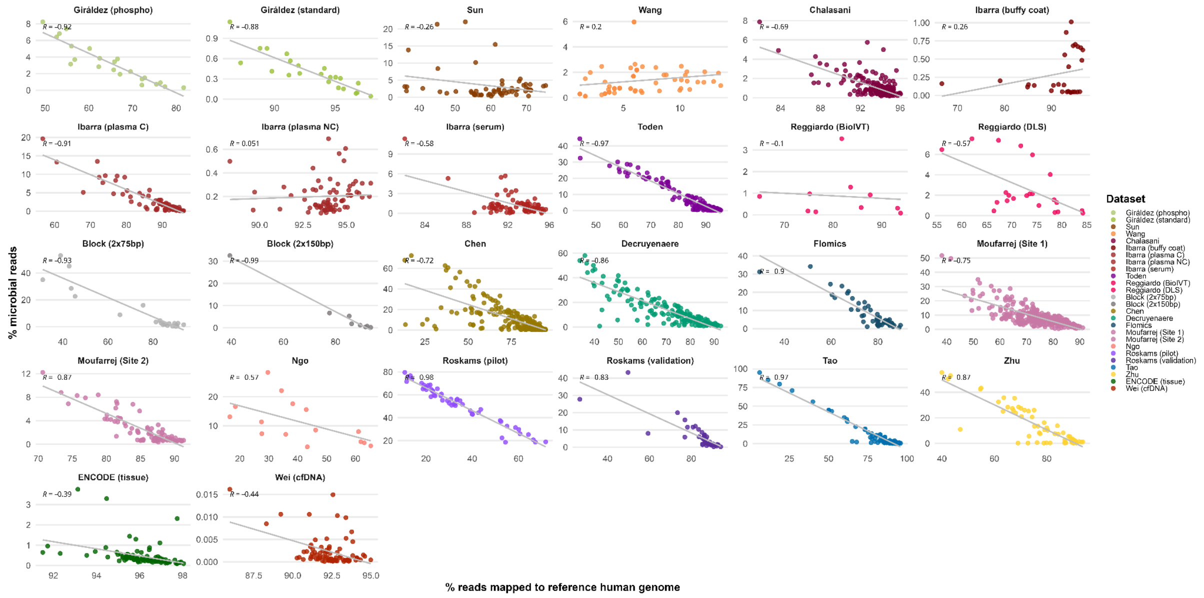

### Supplementary figure 4

**Scatterplot showing the concordance between NG80 and other diversity metrics. (A)** Shannon entropy, **(B)** and Gini index. Each dot corresponds to a sample, colored by dataset. Pearson correlation (R) is reported in the top left corner of each plot.

A

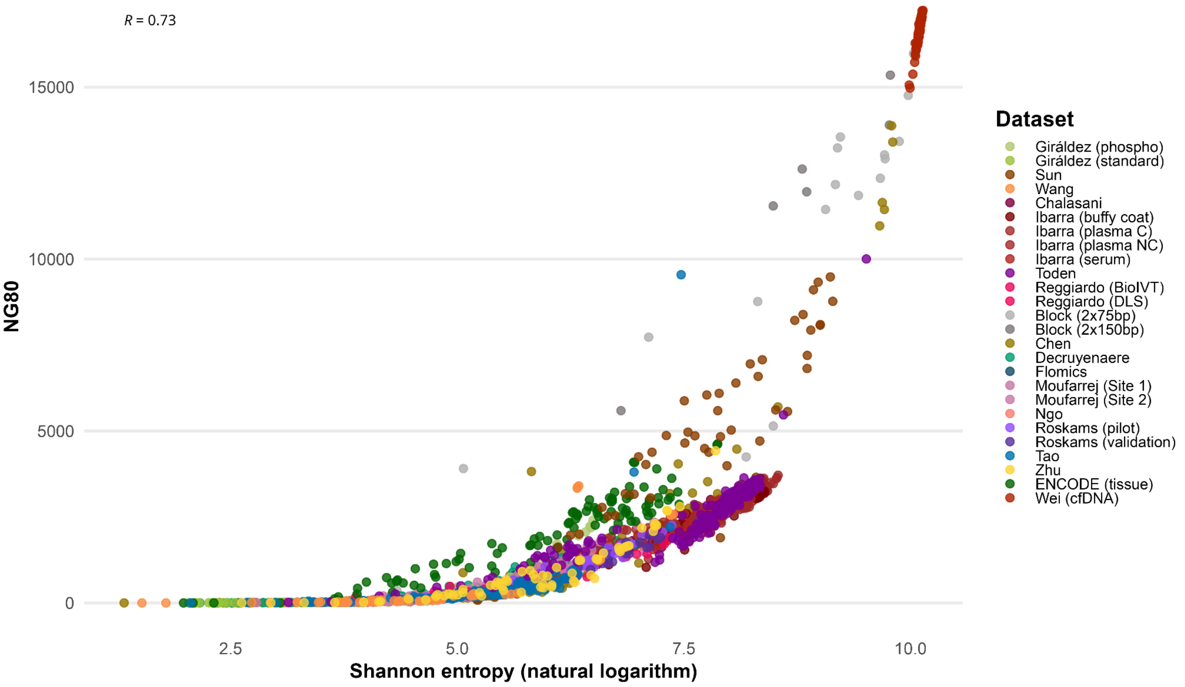

B

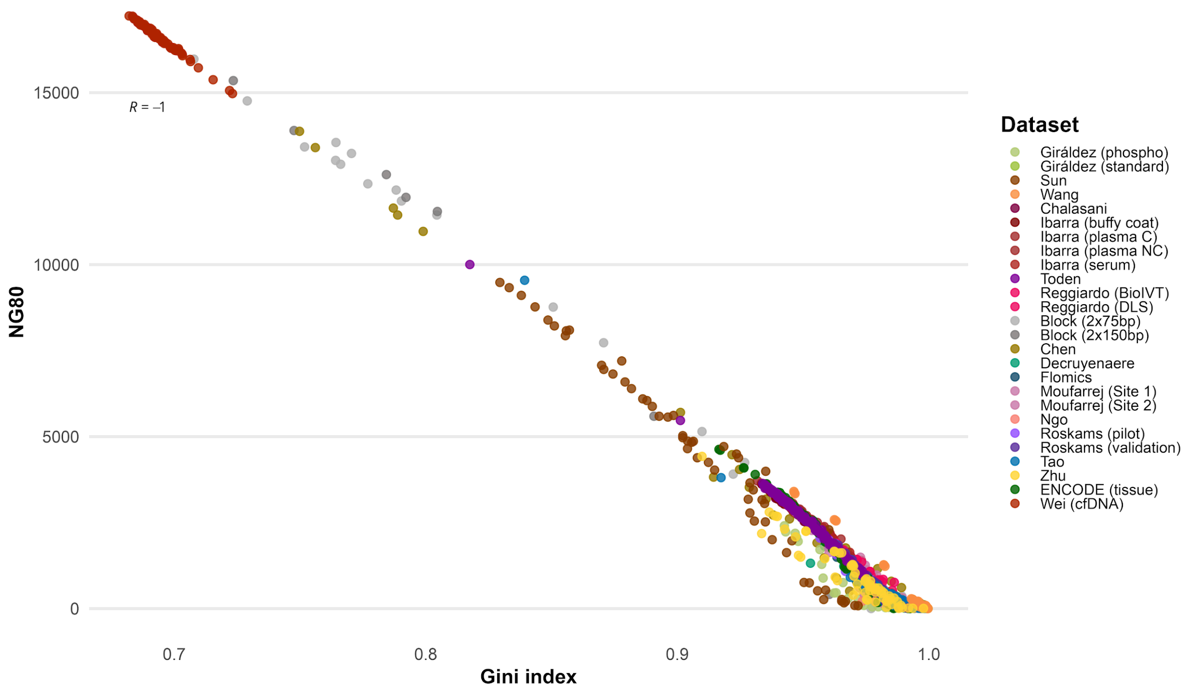

R = -1
15000
10000
5000
0
NG80
0.7
0.8
0.9
1.0
Gini index
Dataset
Giráldez (phospho)
Giráldez (standard)
Sun
Wang
Chalasani
Ibarra (buffy coat)
Ibarra (plasma C)
Ibarra (plasma NC)
Ibarra (serum)
Toden
Reggiardo (BioIVT)
Reggiardo (DLS)
Block (2x75bp)
Block (2x150bp)
Chen
Decruyenaere
Flomics
Moufarrej (Site 1)
Moufarrej (Site 2)
Ngo
Roskams (pilot)
Roskams (validation)
Tao
Zhu
ENCODE (tissue)
Wei (cfDNA)

### Supplementary figure 5

**Distribution of RNA cell-type contribution prediction across gDNA-free datasets**. Major cell types (MCTs, namely, cell types with at least 10% of predicted contribution in at least the 10% of the sample) were selected from each dataset and then collapsed in a unique list (i.e. erythrocyte, platelet, monocyte, b cell, neutrophil, macrophage). Both Chen and Zhu datasets were split in two batches, based on differences in cell type contribution (see main text).

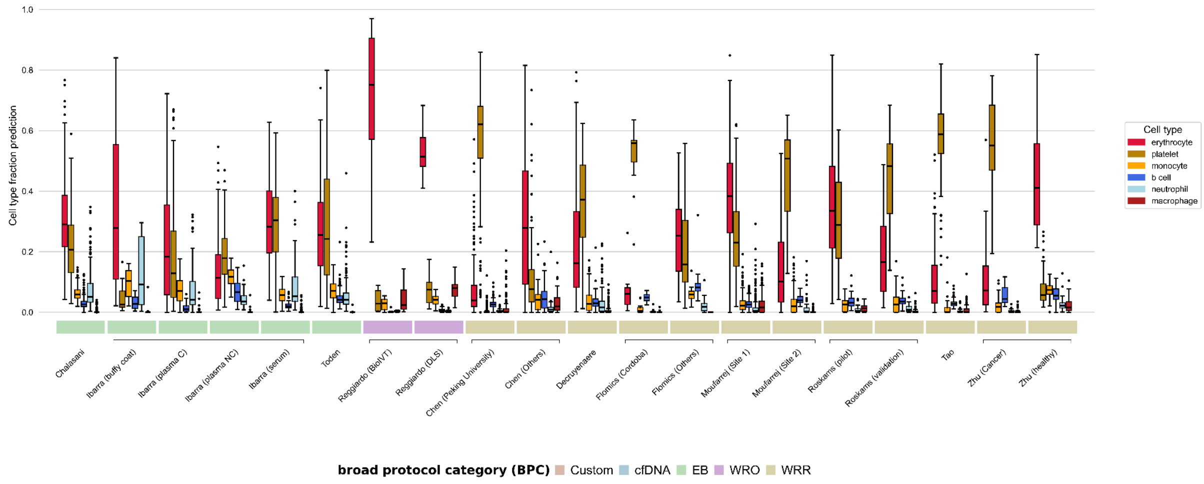

### Supplementary figure 6

**Clustered heatmaps showing the predicted cell type contribution to the cell-free transcriptome across datasets**. Only major cell types (MCTs) are represented, on the y axis. Heatmap cells are colored according to the corresponding predicted cell type contribution to the cfRNA library (x axis), in relative abundance. Each library is annotated with its corresponding relevant metadata. **(A)** Flomics, **(B)** Chalasani, **(C)** Ibarra, **(D)** Moufarrej, **(E)** Reggiardo, **(F)** Roskams-Hieter, **(G)** Tao, **(H)** Toden, **(I)** Zhu.

A

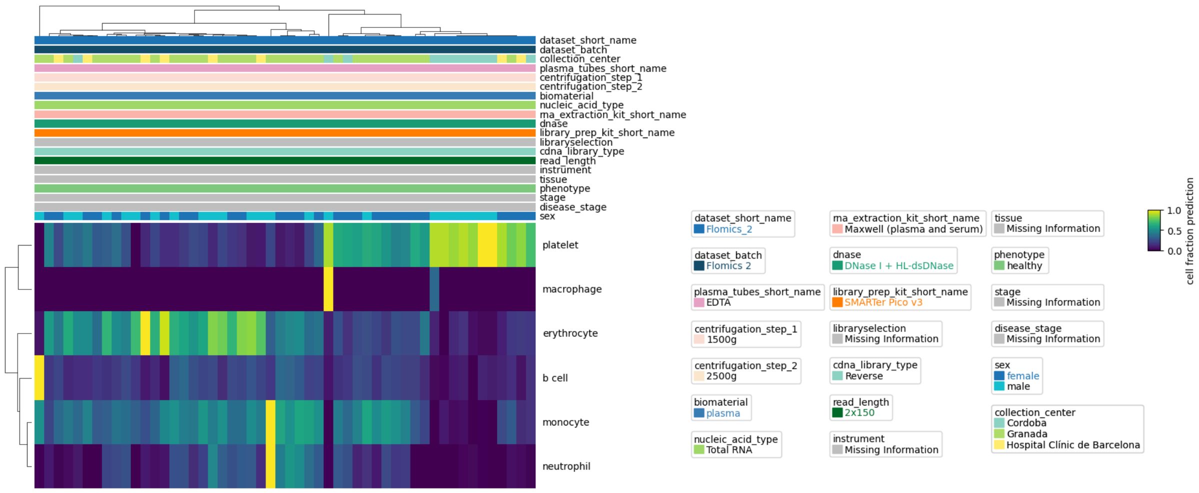


B

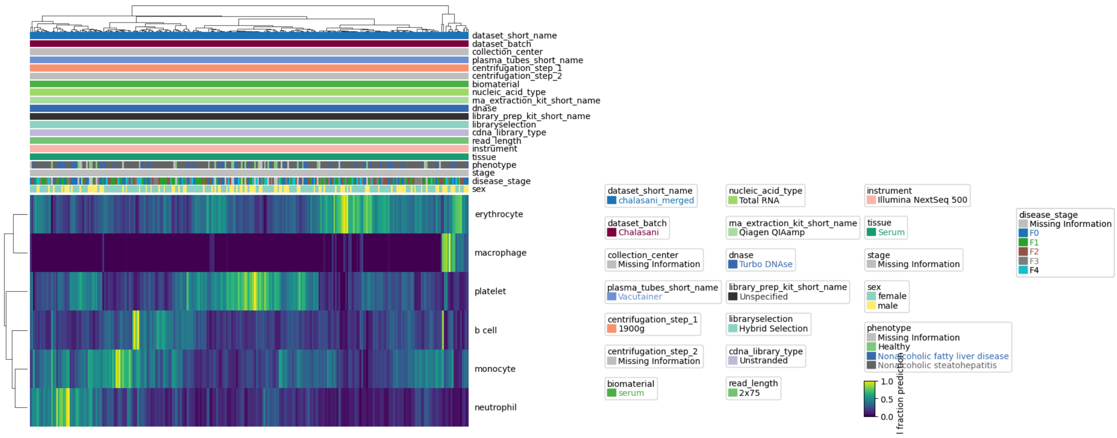

C

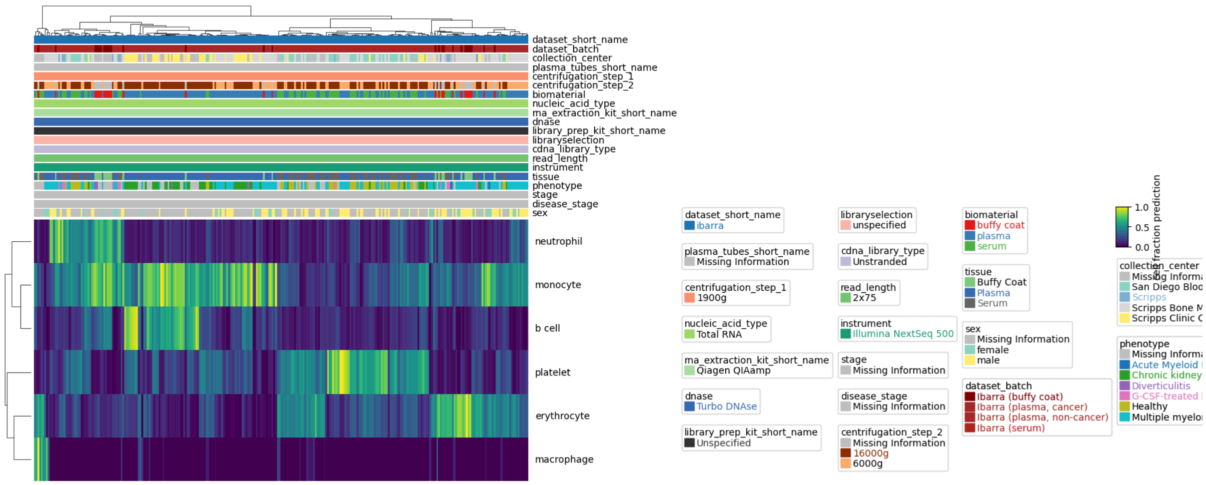

dataset_short_name
dataset_batch
collection_center
plasma_tubes_short_name
centrifugation_step_1
centrifugation_step_2
biomaterial
nucleic_acid_type
rna_extraction_kit_short_name
dnase
library_prep_kit_short_name
libraryselection
cdna_library_type
read_length
instrument
tissue
phenotype
stage
disease_stage
sex
neutrophil
monocyte
b cell
platelet
erythrocyte
macrophage
dataset_short_name: ibarra
plasma_tubes_short_name: Missing Information
centrifugation_step_1: 1900g
nucleic_acid_type: Total RNA
rna_extraction_kit_short_name: Qiagen QIAamp
dnase: Turbo DNAse
library_prep_kit_short_name: Unspecified
libraryselection: unspecified
cdna_library_type: Unstranded
read_length: 2x75
instrument: Illumina NextSeq 500
stage: Missing Information
disease_stage: Missing Information
centrifugation_step_2: Missing Information, 16000g, 6000g
biomaterial: buffy coat, plasma, serum
tissue: Buffy Coat, Plasma, Serum
sex: Missing Information, female, male
dataset_batch: Ibarra (buffy coat), Ibarra (plasma, cancer), Ibarra (plasma, non-cancer), Ibarra (serum)
collection_center
phenotype
Healthy
1.0
0.5
0.0
cell fraction prediction


D

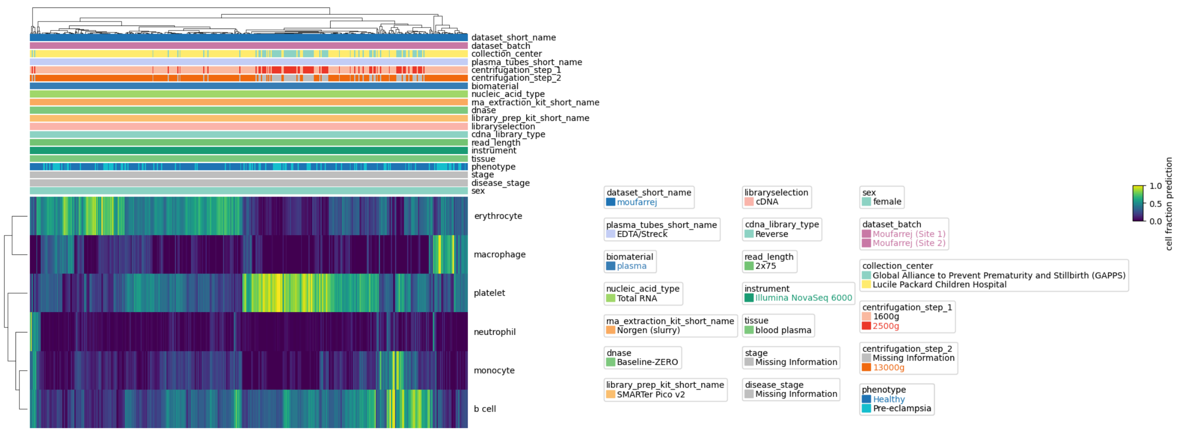

dataset_short_name
dataset_batch
collection_center
plasma_tubes_short_name
centrifugation_step_1
centrifugation_step_2
biomaterial
nucleic_acid_type
rna_extraction_kit_short_name
dnase
library_prep_kit_short_name
libraryselection
cdna_library_type
read_length
instrument
tissue
phenotype
stage
disease_stage
sex
erythrocyte
macrophage
platelet
neutrophil
monocyte
b cell
dataset_short_name: moufarrej
plasma_tubes_short_name: EDTA/Streck
biomaterial: plasma
nucleic_acid_type: Total RNA
rna_extraction_kit_short_name: Norgen (slurry)
dnase: Baseline-ZERO
library_prep_kit_short_name: SMARTer Pico v2
libraryselection: cDNA
cdna_library_type: Reverse
read_length: 2x75
instrument: Illumina NovaSeq 6000
tissue: blood plasma
stage: Missing Information
disease_stage: Missing Information
sex: female
dataset_batch: Moufarrej (Site 1), Moufarrej (Site 2)
collection_center: Global Alliance to Prevent Prematurity and Stillbirth (GAPPS), Lucile Packard Children Hospital
centrifugation_step_1: 1600g, 2500g
centrifugation_step_2: Missing Information, 13000g
phenotype: Healthy, Pre-eclampsia
1.0
0.5
0.0
cell fraction prediction


E

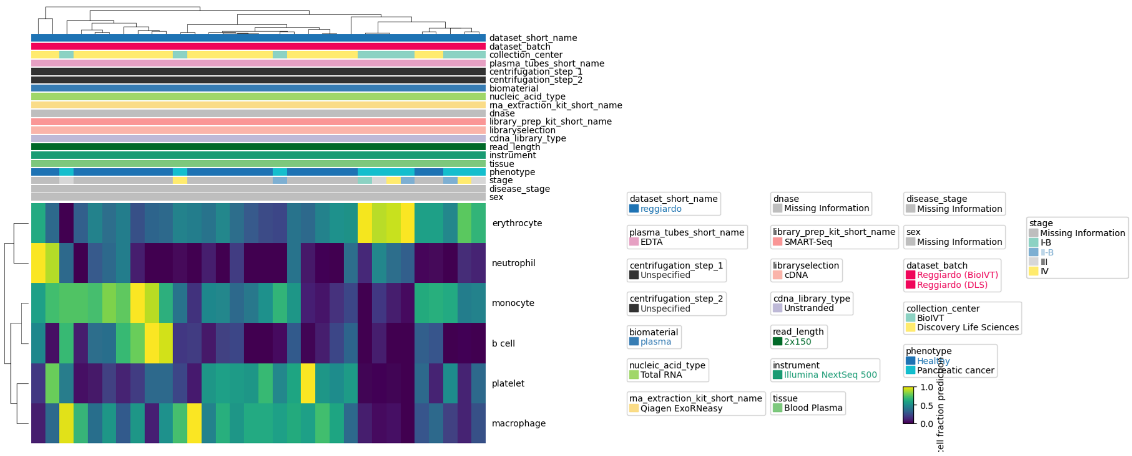

dataset_short_name
dataset_batch
collection_center
plasma_tubes_short_name
centrifugation_step_1
centrifugation_step_2
biomaterial
nucleic_acid_type
rna_extraction_kit_short_name
dnase
library_prep_kit_short_name
libraryselection
cdna_library_type
read_length
instrument
tissue
phenotype
stage
disease_stage
sex
erythrocyte
neutrophil
monocyte
b cell
platelet
macrophage
dataset_short_name: reggiardo
plasma_tubes_short_name: EDTA
centrifugation_step_1: Unspecified
centrifugation_step_2: Unspecified
biomaterial: plasma
nucleic_acid_type: Total RNA
rna_extraction_kit_short_name: Qiagen ExoRNeasy
dnase: Missing Information
library_prep_kit_short_name: SMART-Seq
libraryselection: cDNA
cdna_library_type: Unstranded
read_length: 2x150
instrument: Illumina NextSeq 500
tissue: Blood Plasma
disease_stage: Missing Information
sex: Missing Information
dataset_batch: Reggiardo (BioIVT), Reggiardo (DLS)
collection_center: BioIVT, Discovery Life Sciences
phenotype: Healthy, Pancreatic cancer
stage: Missing Information, I-B, II-B, III, IV
1.0
0.5
0.0

F

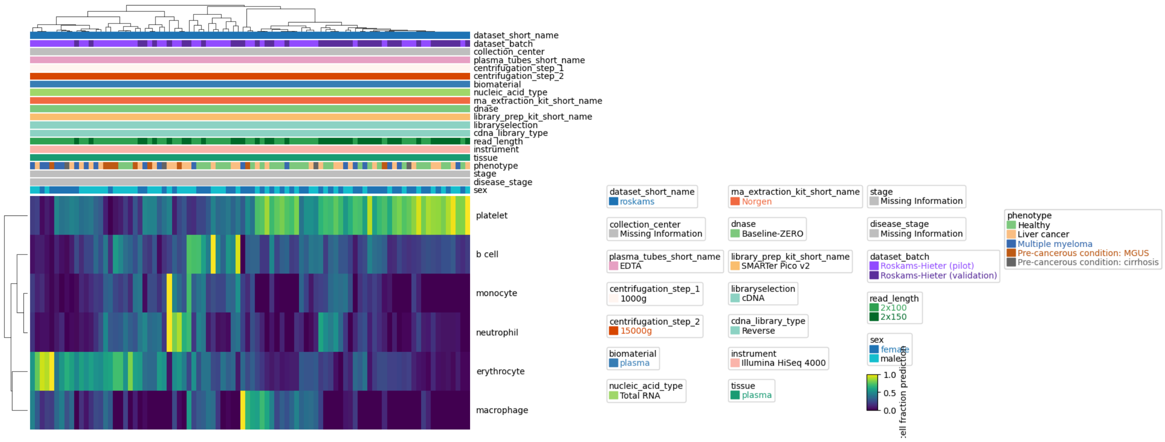
dataset_short_name
dataset_batch
collection_center
plasma_tubes_short_name
centrifugation_step_1
centrifugation_step_2
biomaterial
nucleic_acid_type
rna_extraction_kit_short_name
dnase
library_prep_kit_short_name
libraryselection
cdna_library_type
read_length
instrument
tissue
phenotype
stage
disease_stage
sex
platelet
b cell
monocyte
neutrophil
erythrocyte
macrophage
dataset_short_name roskams
collection_center Missing Information
plasma_tubes_short_name EDTA
centrifugation_step_1 1000g
centrifugation_step_2 15000g
biomaterial plasma
nucleic_acid_type Total RNA
rna_extraction_kit_short_name Norgen
dnase Baseline-ZERO
library_prep_kit_short_name SMARTer Pico v2
libraryselection cDNA
cdna_library_type Reverse
instrument Illumina HiSeq 4000
tissue plasma
stage Missing Information
disease_stage Missing Information
dataset_batch Roskams-Hieter (pilot) Roskams-Hieter (validation)
read_length 2x100 2x150
sex female male
1.0
0.5
0.0
phenotype Healthy Liver cancer Multiple myeloma Pre-cancerous condition: MGUS Pre-cancerous condition: cirrhosis

G

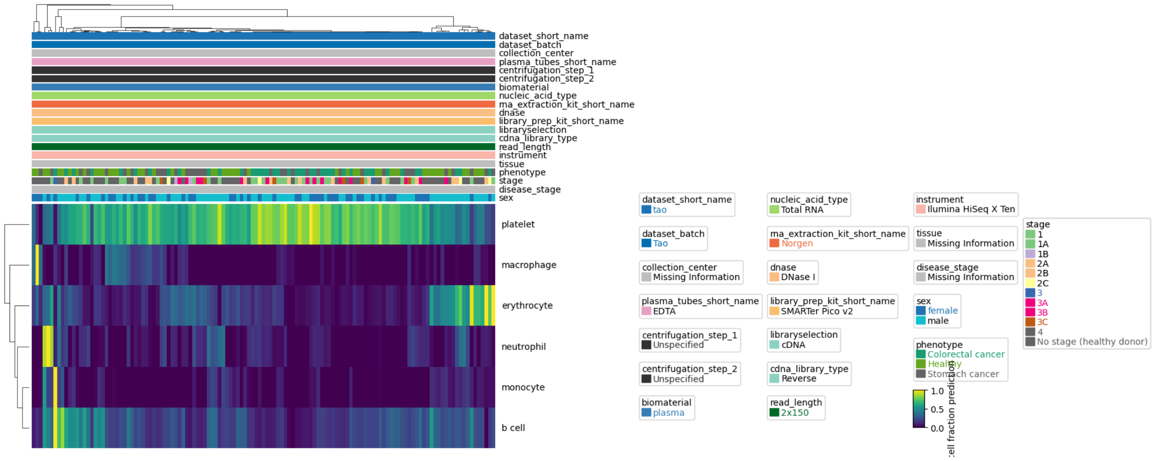
dataset_short_name
dataset_batch
collection_center
plasma_tubes_short_name
centrifugation_step_1
centrifugation_step_2
biomaterial
nucleic_acid_type
rna_extraction_kit_short_name
dnase
library_prep_kit_short_name
libraryselection
cdna_library_type
read_length
instrument
tissue
phenotype
stage
disease_stage
sex
platelet
macrophage
erythrocyte
neutrophil
monocyte
b cell
dataset_short_name tao
dataset_batch Tao
collection_center Missing Information
plasma_tubes_short_name EDTA
centrifugation_step_1 Unspecified
centrifugation_step_2 Unspecified
biomaterial plasma
nucleic_acid_type Total RNA
rna_extraction_kit_short_name Norgen
dnase DNase I
library_prep_kit_short_name SMARTer Pico v2
libraryselection cDNA
cdna_library_type Reverse
read_length 2x150
instrument Ilumina HiSeq X Ten
tissue Missing Information
disease_stage Missing Information
sex female male
phenotype Colorectal cancer Healthy Stomach cancer
1.0
0.5
0.0
stage 1 1A 1B 2A 2B 2C 3 3A 3B 3C 4 No stage (healthy donor)

H

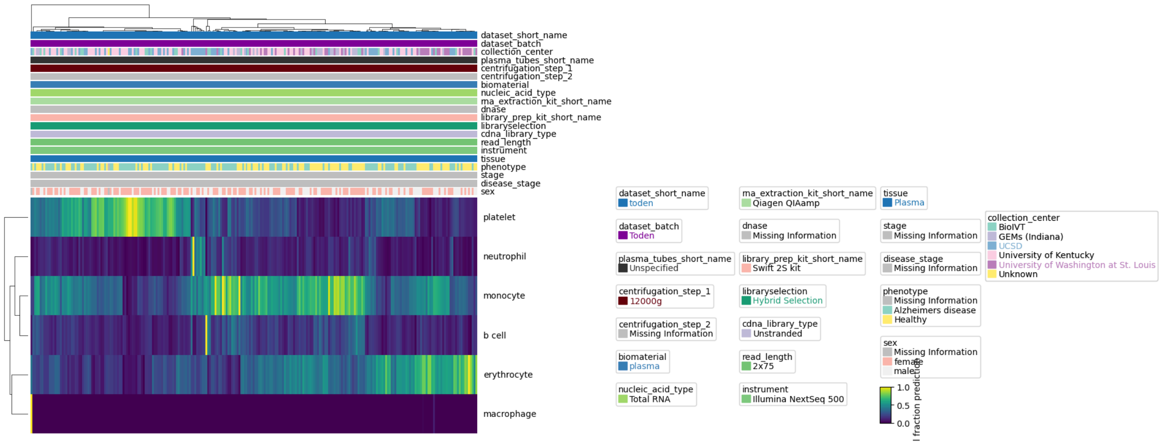
dataset_short_name
dataset_batch
collection_center
plasma_tubes_short_name
centrifugation_step_1
centrifugation_step_2
biomaterial
nucleic_acid_type
rna_extraction_kit_short_name
dnase
library_prep_kit_short_name
libraryselection
cdna_library_type
read_length
instrument
tissue
phenotype
stage
disease_stage
sex
platelet
neutrophil
monocyte
b cell
erythrocyte
macrophage
dataset_short_name toden
dataset_batch Toden
plasma_tubes_short_name Unspecified
centrifugation_step_1 12000g
centrifugation_step_2 Missing Information
biomaterial plasma
nucleic_acid_type Total RNA
rna_extraction_kit_short_name Qiagen QIAamp
dnase Missing Information
library_prep_kit_short_name Swift 2S kit
libraryselection Hybrid Selection
cdna_library_type Unstranded
read_length 2x75
instrument Illumina NextSeq 500
tissue Plasma
stage Missing Information
disease_stage Missing Information
phenotype Missing Information Alzheimers disease Healthy
sex Missing Information female male
1.0
0.5
0.0
collection_center BioIVT GEMs (Indiana) UCSD University of Kentucky University of Washington at St. Louis Unknown

I

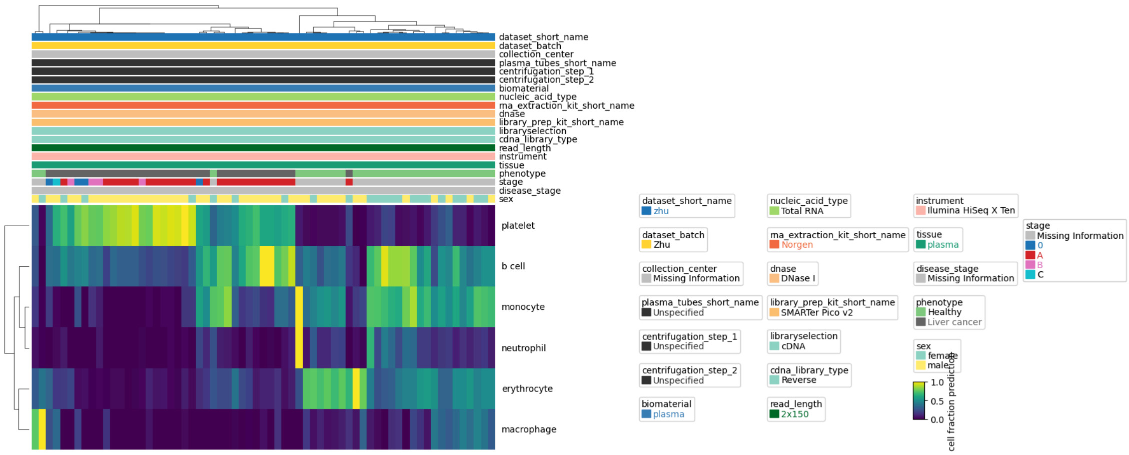
dataset_short_name
dataset_batch
collection_center
plasma_tubes_short_name
centrifugation_step_1
centrifugation_step_2
biomaterial
nucleic_acid_type
rna_extraction_kit_short_name
dnase
library_prep_kit_short_name
libraryselection
cdna_library_type
read_length
instrument
tissue
phenotype
stage
disease_stage
sex
platelet
b cell
monocyte
neutrophil
erythrocyte
macrophage
dataset_short_name
zhu
dataset_batch
Zhu
collection_center
Missing Information
plasma_tubes_short_name
Unspecified
centrifugation_step_1
Unspecified
centrifugation_step_2
Unspecified
biomaterial
plasma
nucleic_acid_type
Total RNA
rna_extraction_kit_short_name
Norgen
dnase
DNase I
library_prep_kit_short_name
SMARTer Pico v2
libraryselection
cDNA
cdna_library_type
Reverse
read_length
2x150
instrument
Ilumina HiSeq X Ten
tissue
plasma
disease_stage
Missing Information
phenotype
Healthy
Liver cancer
sex
female
male
stage
Missing Information
0
A
B
C
1.0
0.5
0.0
cell fraction prediction

### Supplementary figure 7

**Dimensionality reduction of gene abundance profiles**. Scatter-plot visualizations of sequencing libraries in reduced gene abundance space using t-distributed stochastic neighbor embedding (t-SNE) and principal component analysis (PCA). Each point represents one sequencing library. (**A**) t-SNE plot colored by dataset/batch-label, including all datasets in analysis. **(B)** PCA with healthy controls only, colored by dataset and batch. **(C)** PCA showing sample clustering before (top) and after (bottom) linear-model residualization. Samples are colored according to their original dataset (right) and patient phenotype (left). (D-G) PCA including sequencing libraries from (**D**) Chen, **(E)** Flomics, (**F**) Moufarrej (**G**) Zhu. Dots are colored by factor driving batch effect (collection center, left) and relative abundance of platelet RNA (right). **(H)** gDNA-free samples. **(I)** gDNA-free datasets, colored by dataset/batch label.

A

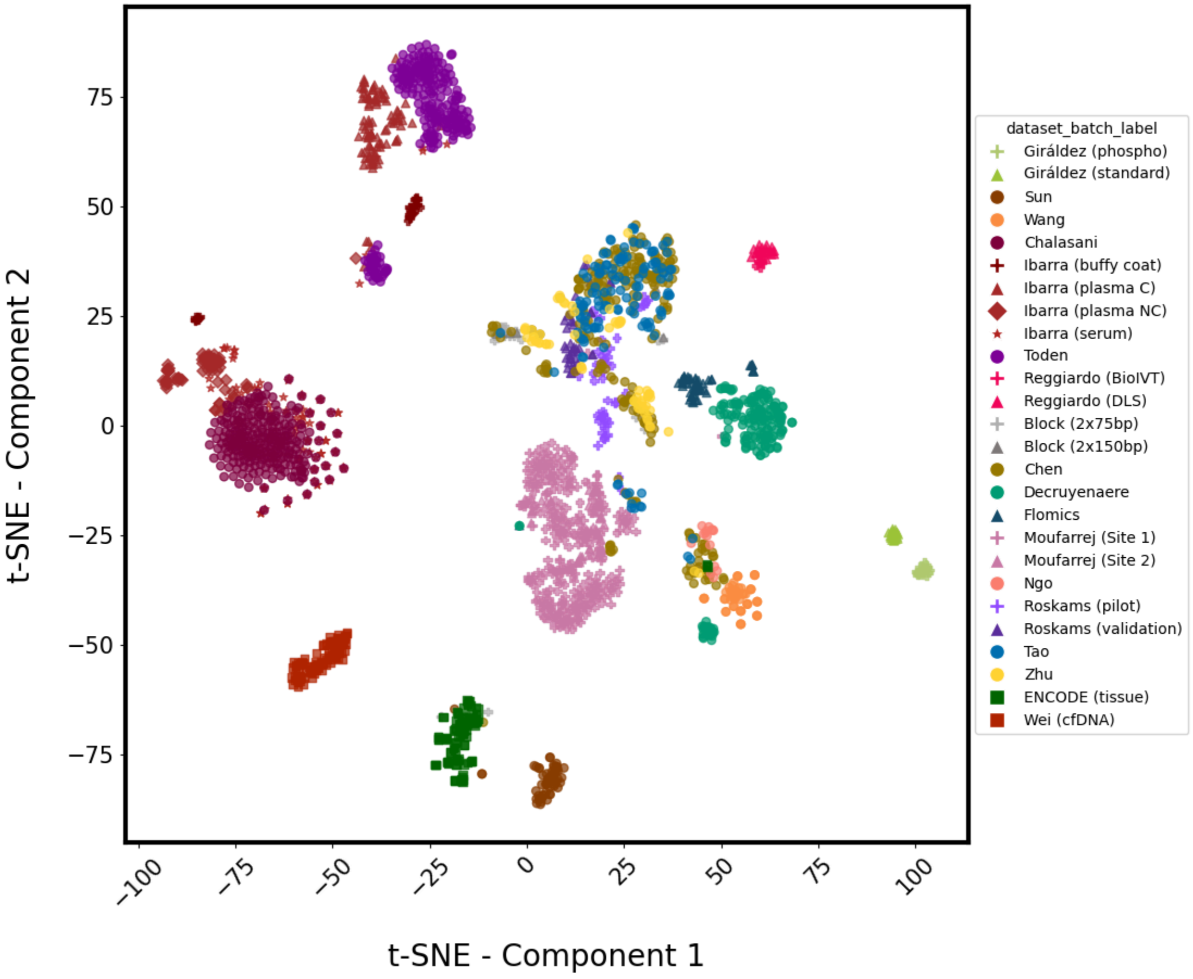

B

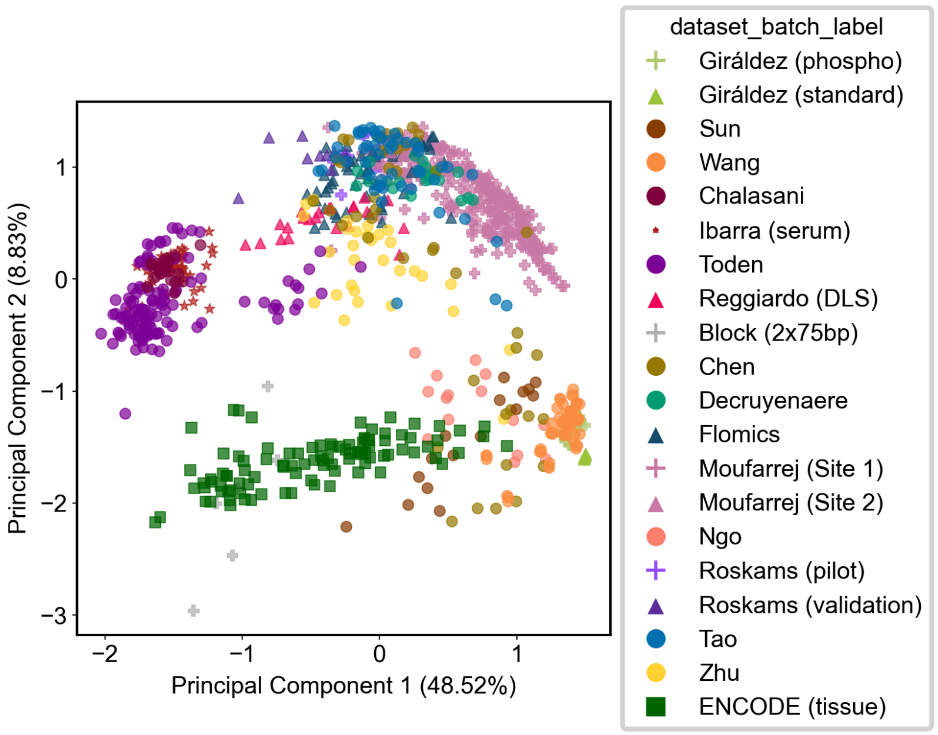
Principal Component 2 (8.83%)
Principal Component 1 (48.52%)
1
0
−1
−2
−3
−2
−1
0
1
dataset_batch_label
Giráldez (phospho)
Giráldez (standard)
Sun
Wang
Chalasani
Ibarra (serum)
Toden
Reggiardo (DLS)
Block (2x75bp)
Chen
Decruyenaere
Flomics
Moufarrej (Site 1)
Moufarrej (Site 2)
Ngo
Roskams (pilot)
Roskams (validation)
Tao
Zhu
ENCODE (tissue)

C

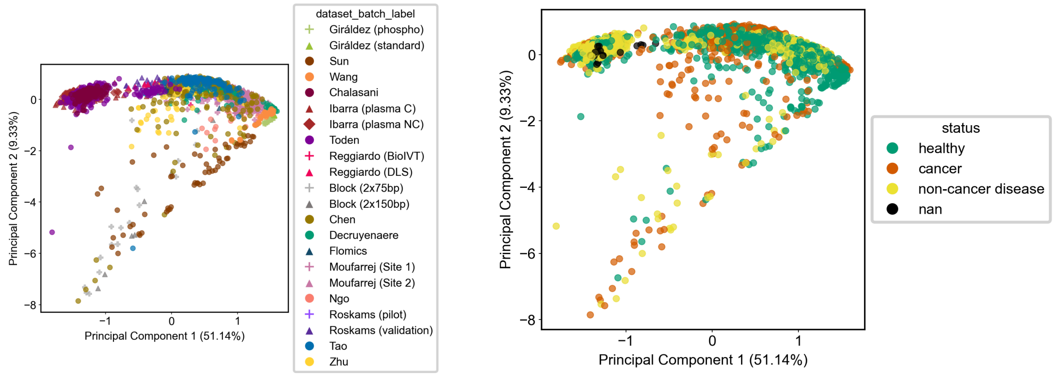
Principal Component 2 (9.33%)
Principal Component 1 (51.14%)
0
−2
−4
−6
−8
−1
0
1
dataset_batch_label
Giráldez (phospho)
Giráldez (standard)
Sun
Wang
Chalasani
Ibarra (plasma C)
Ibarra (plasma NC)
Toden
Reggiardo (BioIVT)
Reggiardo (DLS)
Block (2x75bp)
Block (2x150bp)
Chen
Decruyenaere
Flomics
Moufarrej (Site 1)
Moufarrej (Site 2)
Ngo
Roskams (pilot)
Roskams (validation)
Tao
Zhu
status
healthy
cancer
non-cancer disease
nan

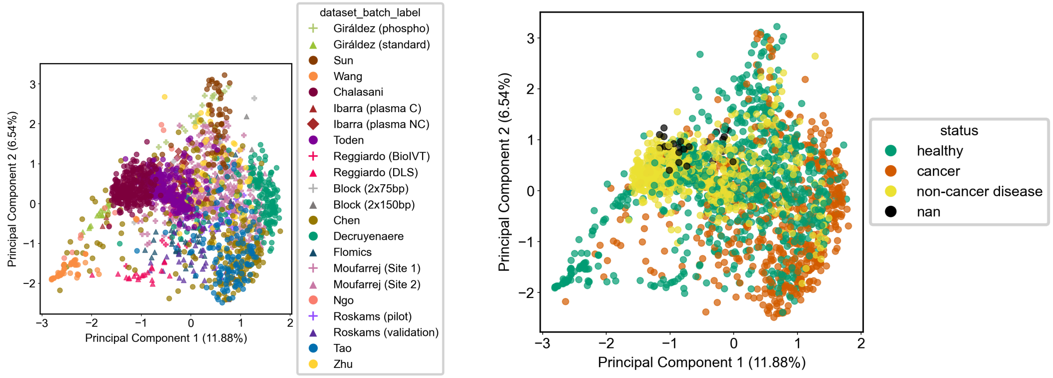
dataset_batch_label
Giráldez (phospho)
Giráldez (standard)
Sun
Wang
Chalasani
Ibarra (plasma C)
Ibarra (plasma NC)
Toden
Reggiardo (BioIVT)
Reggiardo (DLS)
Block (2x75bp)
Block (2x150bp)
Chen
Decruyenaere
Flomics
Moufarrej (Site 1)
Moufarrej (Site 2)
Ngo
Roskams (pilot)
Roskams (validation)
Tao
Zhu
Principal Component 1 (11.88%)
Principal Component 2 (6.54%)
status
healthy
cancer
non-cancer disease
nan

D

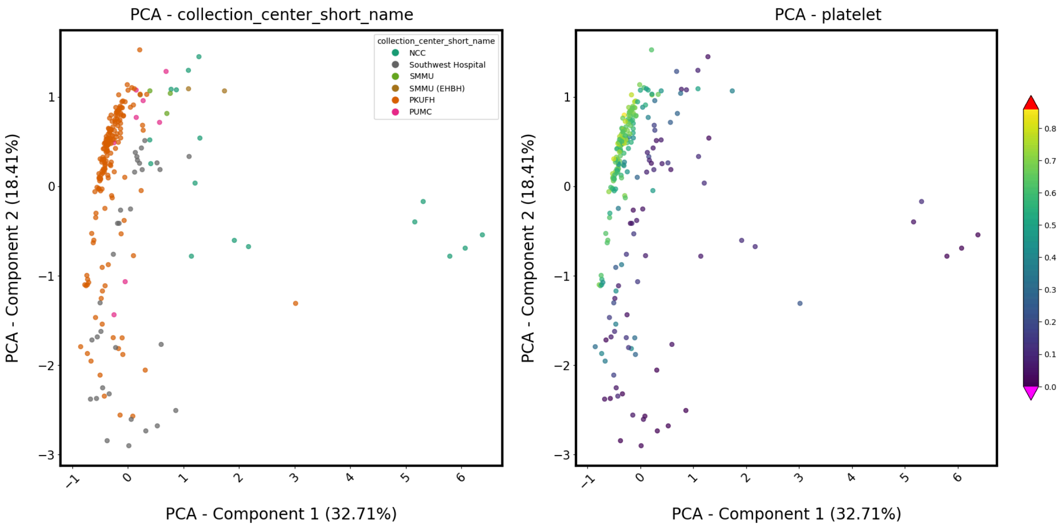
PCA - collection_center_short_name
collection_center_short_name
NCC
Southwest Hospital
SMMU
SMMU (EHBH)
PKUFH
PUMC
PCA - platelet
PCA - Component 1 (32.71%)
PCA - Component 2 (18.41%)

E

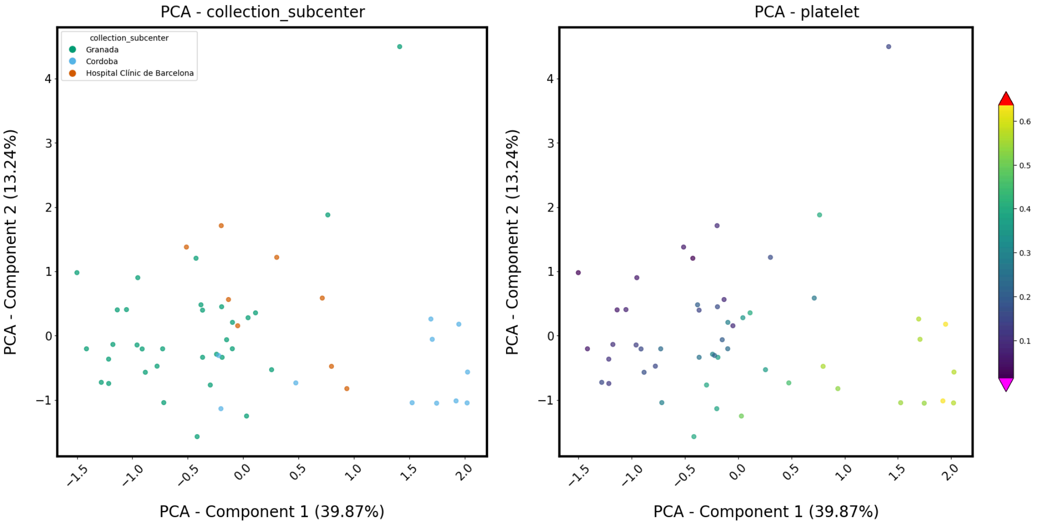
PCA - collection_subcenter
collection_subcenter
Granada
Cordoba
Hospital Clínic de Barcelona
PCA - Component 2 (13.24%)
PCA - Component 1 (39.87%)
PCA - platelet
PCA - Component 2 (13.24%)
PCA - Component 1 (39.87%)

F

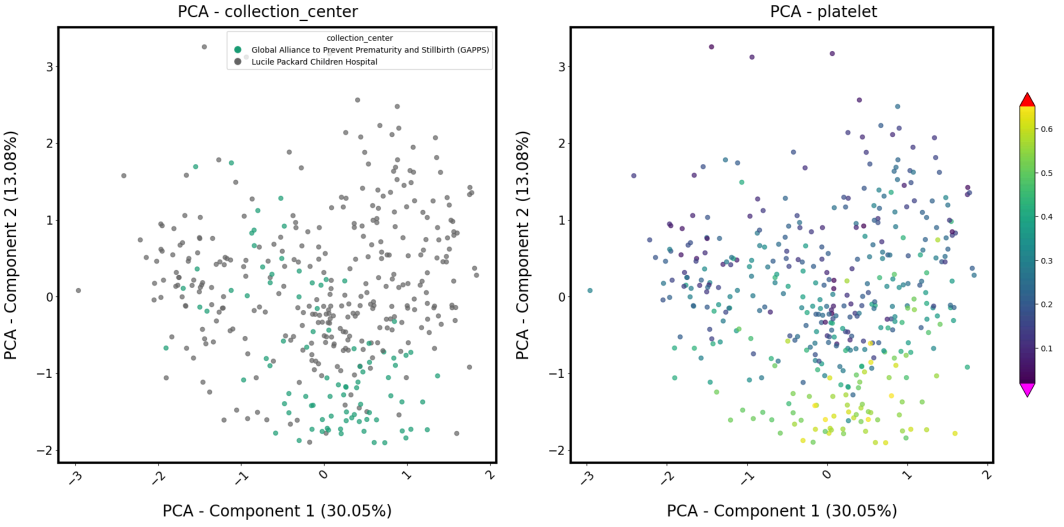
PCA - collection_center
collection_center
Global Alliance to Prevent Prematurity and Stillbirth (GAPPS)
Lucile Packard Children Hospital
PCA - Component 2 (13.08%)
PCA - Component 1 (30.05%)
PCA - platelet
PCA - Component 2 (13.08%)
PCA - Component 1 (30.05%)

G

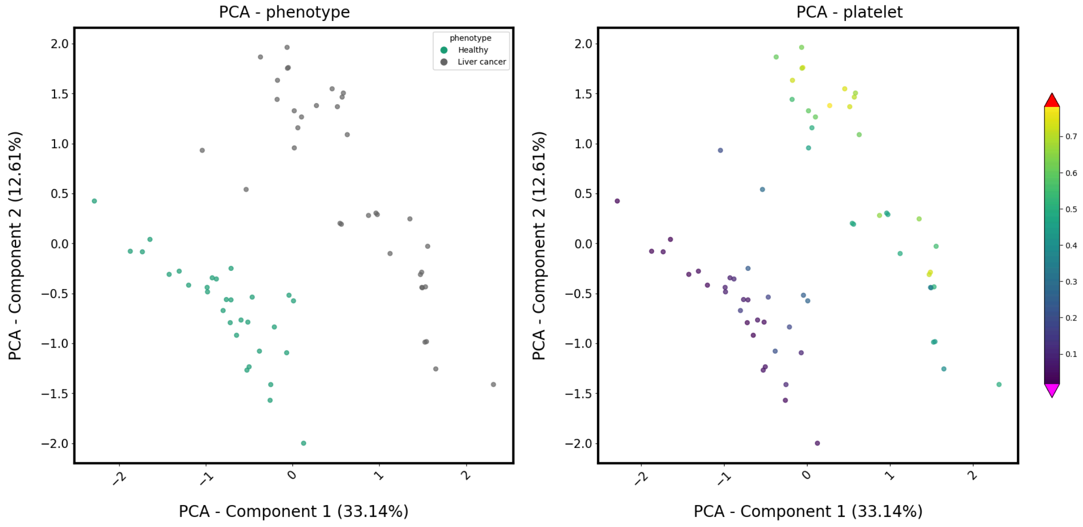

PCA - phenotype
PCA - platelet
phenotype
Healthy
Liver cancer
PCA - Component 1 (33.14%)
PCA - Component 2 (12.61%)


H

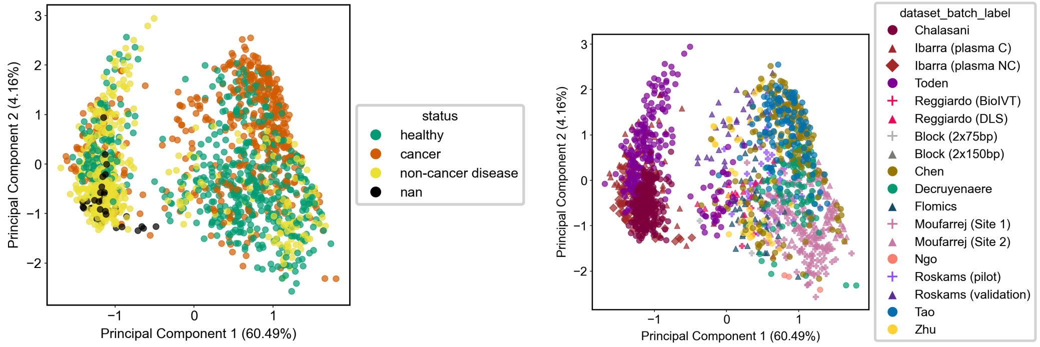

status
healthy
cancer
non-cancer disease
nan
Principal Component 1 (60.49%)
Principal Component 2 (4.16%)
dataset_batch_label
Chalasani
Ibarra (plasma C)
Ibarra (plasma NC)
Toden
Reggiardo (BioIVT)
Reggiardo (DLS)
Block (2x75bp)
Block (2x150bp)
Chen
Decruyenaere
Flomics
Moufarrej (Site 1)
Moufarrej (Site 2)
Ngo
Roskams (pilot)
Roskams (validation)
Tao
Zhu

**I**

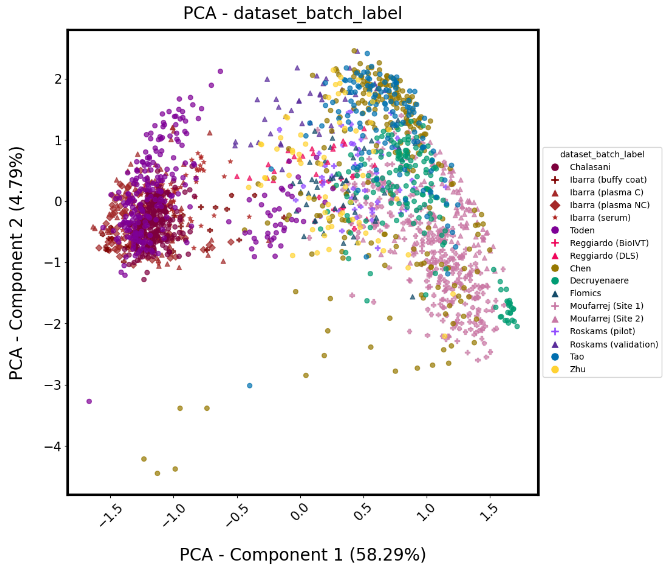
PCA - dataset_batch_label
PCA - Component 2 (4.79%)
PCA - Component 1 (58.29%)
dataset_batch_label
Chalasani
Ibarra (buffy coat)
Ibarra (plasma C)
Ibarra (plasma NC)
Ibarra (serum)
Toden
Reggiardo (BioIVT)
Reggiardo (DLS)
Chen
Decruyenaere
Flomics
Moufarrej (Site 1)
Moufarrej (Site 2)
Roskams (pilot)
Roskams (validation)
Tao
Zhu

### Supplementary figure 8

**Concordance between gene abundance profiles obtained with filtered (human-only) and unfiltered reads as a function of effective fragment length**. **(A)** Scatterplot illustrating the relationship between the effective fragment length (EFL, X axis) and the Pearson correlation coefficient (r, Y axis) for gene-level quantifications. Correlations were calculated on log10(raw counts) + 1) between the "unfiltered" and "human-only" (filtered) reads-based gene quantifications for each biological sample. Individual data points represent single samples, color-coded by their respective dataset. A LOESS regression curve (solid black line) is fitted to the data to highlight the non-linear trend, revealing a distinct inflection point in quantification consistency at > 100bp. **(B)** Scatterplots showing the lack of association between fraction of human reads ("percent_human_reads (%)", X axis) and Pearson correlation coefficient between gene quantifications obtained with filtered (human-only) and unfiltered reads ("Pearson R", Y axis). Each subplot corresponds to a dataset/batch. Each dot represents a sample, and is colored according to its average EFL. The Pearson (r) correlation between the X and Y variables and sample size (n) are reported for each dataset/batch in the bottom right corner of the corresponding subplot.

A

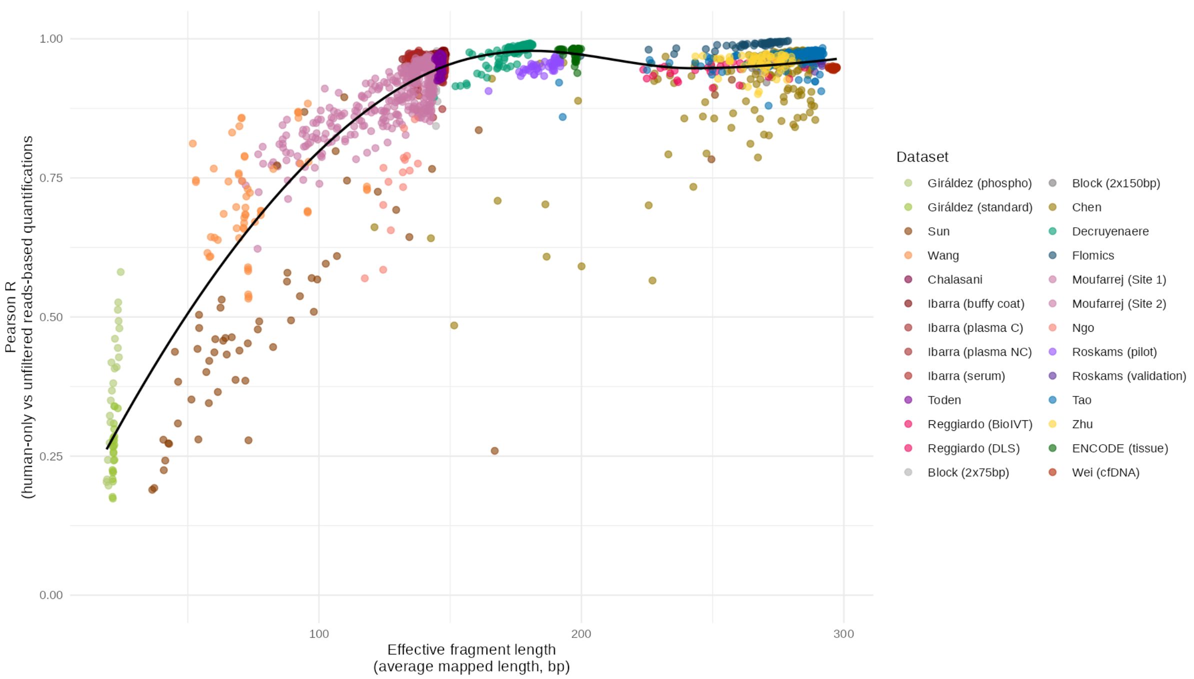
1.00
0.75
0.50
0.25
0.00
Pearson R
(human-only vs unfiltered reads-based quantifications)
100
200
300
Effective fragment length
(average mapped length, bp)
Dataset
Giráldez (phospho)
Giráldez (standard)
Sun
Wang
Chalasani
Ibarra (buffy coat)
Ibarra (plasma C)
Ibarra (plasma NC)
Ibarra (serum)
Toden
Reggiardo (BioIVT)
Reggiardo (DLS)
Block (2x75bp)
Block (2x150bp)
Chen
Decruyenaere
Flomics
Moufarrej (Site 1)
Moufarrej (Site 2)
Ngo
Roskams (pilot)
Roskams (validation)
Tao
Zhu
ENCODE (tissue)
Wei (cfDNA)

B

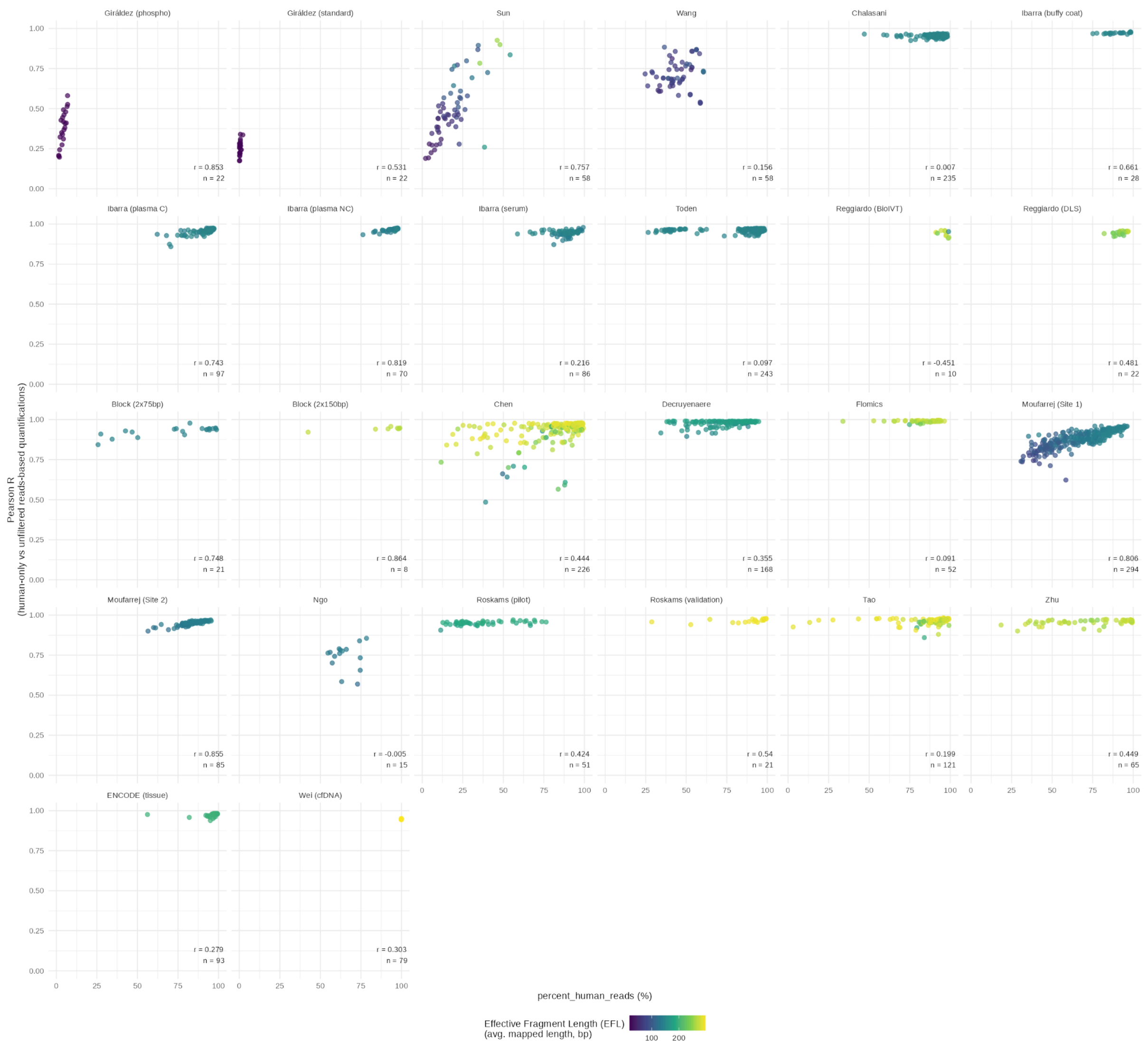

Giráldez (phospho)
r = 0.853
n = 22
Giráldez (standard)
r = 0.531
n = 22
Sun
r = 0.757
n = 58
Wang
r = 0.156
n = 58
Chalasani
r = 0.007
n = 235
Ibarra (buffy coat)
r = 0.661
n = 28
Ibarra (plasma C)
r = 0.743
n = 97
Ibarra (plasma NC)
r = 0.819
n = 70
Ibarra (serum)
r = 0.216
n = 86
Toden
r = 0.097
n = 243
Reggiardo (BioIVT)
r = -0.451
n = 10
Reggiardo (DLS)
r = 0.481
n = 22
Block (2x75bp)
r = 0.748
n = 21
Block (2x150bp)
r = 0.864
n = 8
Chen
r = 0.444
n = 226
Decruyenaere
r = 0.355
n = 168
Flomics
r = 0.091
n = 52
Moufarrej (Site 1)
r = 0.806
n = 294
Moufarrej (Site 2)
r = 0.855
n = 85
Ngo
r = -0.005
n = 15
Roskams (pilot)
r = 0.424
n = 51
Roskams (validation)
r = 0.54
n = 21
Tao
r = 0.199
n = 121
Zhu
r = 0.449
n = 65
ENCODE (tissue)
r = 0.279
n = 93
Wei (cfDNA)
r = 0.303
n = 79
Pearson R
(human-only vs unfiltered reads-based quantifications)
0.00
0.25
0.50
0.75
1.00
0
25
50
75
100
percent_human_reads (%)
Effective Fragment Length (EFL)
(avg. mapped length, bp)
100
200

### Supplementary figure 9

**Sample survival analyses evaluating various quality control thresholds.** The fraction of remaining samples in each dataset is represented at various **(A)** FER, **(B)** FSR, and **(C)** NG80 thresholds.

A

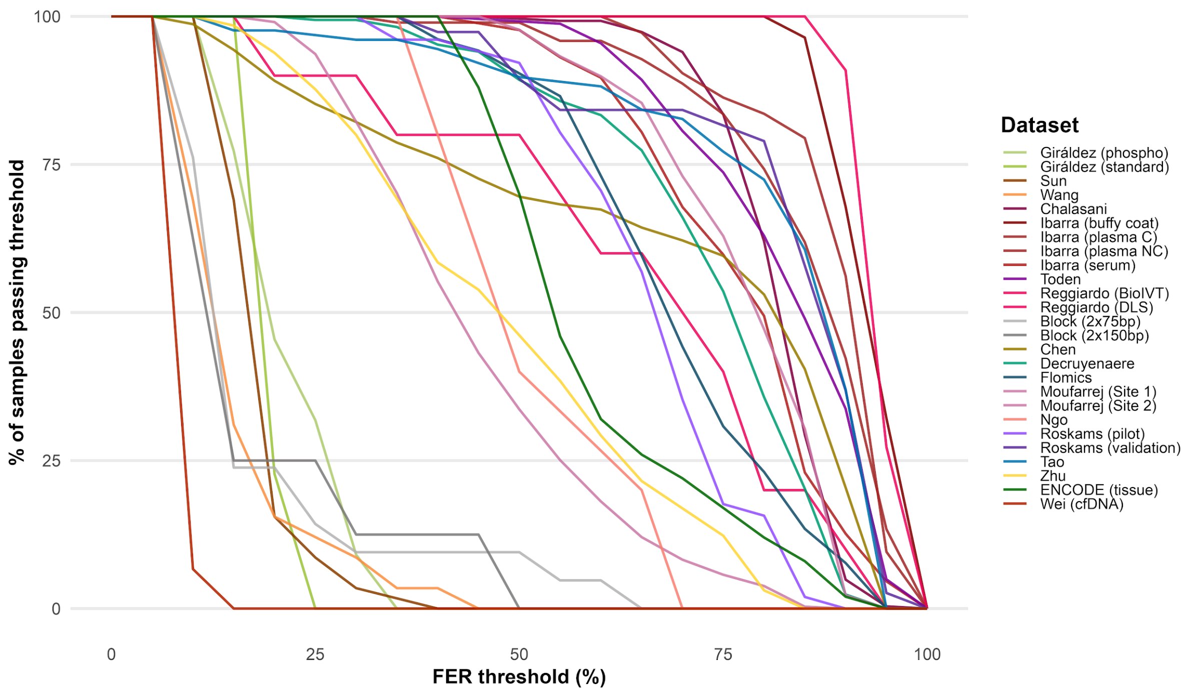

B

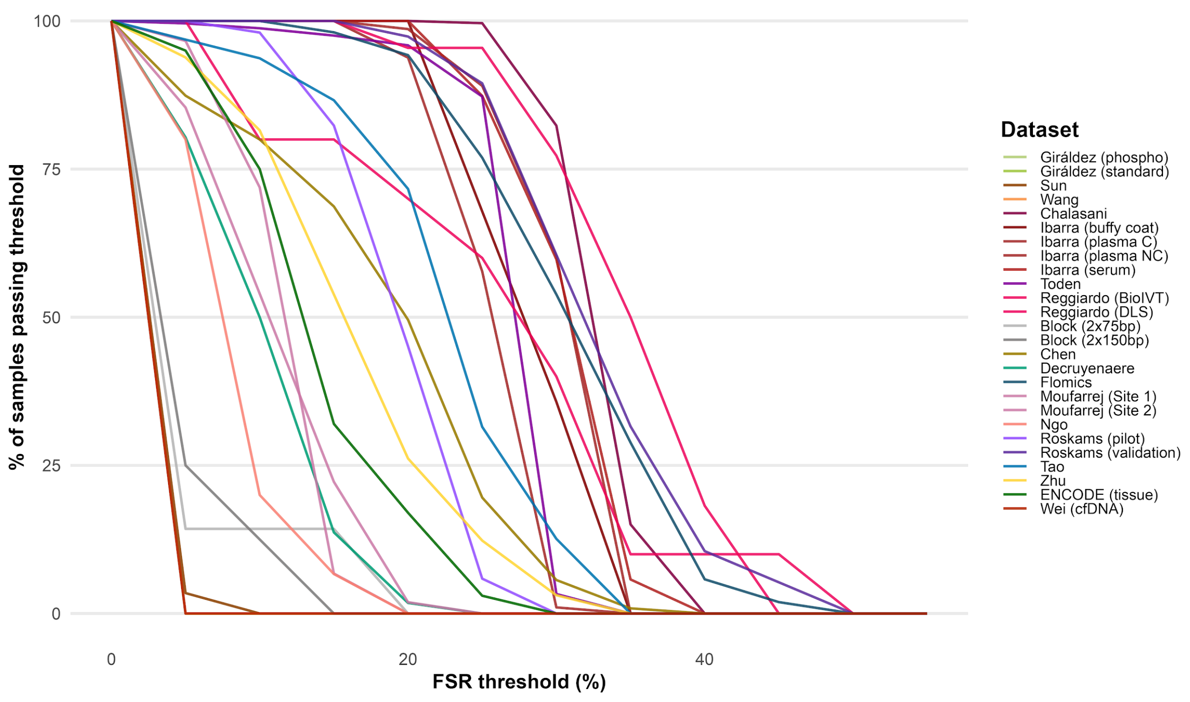
% of samples passing threshold
100
75
50
25
0
0
20
40
FSR threshold (%)
Dataset
Giráldez (phospho)
Giráldez (standard)
Sun
Wang
Chalasani
Ibarra (buffy coat)
Ibarra (plasma C)
Ibarra (plasma NC)
Ibarra (serum)
Toden
Reggiardo (BioIVT)
Reggiardo (DLS)
Block (2x75bp)
Block (2x150bp)
Chen
Decruyenaere
Flomics
Moufarrej (Site 1)
Moufarrej (Site 2)
Ngo
Roskams (pilot)
Roskams (validation)
Tao
Zhu
ENCODE (tissue)
Wei (cfDNA)

C

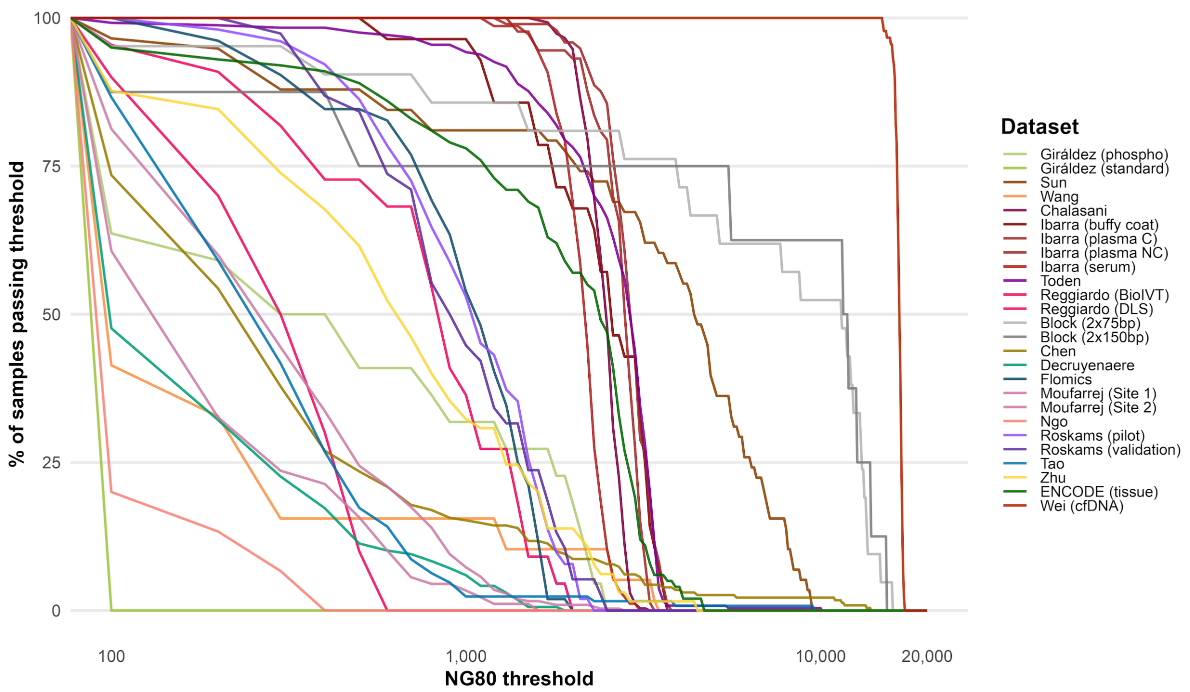
% of samples passing threshold
100
75
50
25
0
100
1,000
10,000
20,000
NG80 threshold
Dataset
Giráldez (phospho)
Giráldez (standard)
Sun
Wang
Chalasani
Ibarra (buffy coat)
Ibarra (plasma C)
Ibarra (plasma NC)
Ibarra (serum)
Toden
Reggiardo (BioIVT)
Reggiardo (DLS)
Block (2x75bp)
Block (2x150bp)
Chen
Decruyenaere
Flomics
Moufarrej (Site 1)
Moufarrej (Site 2)
Ngo
Roskams (pilot)
Roskams (validation)
Tao
Zhu
ENCODE (tissue)
Wei (cfDNA)
